# Variability of the Surface Area of the V1, V2, and V3 Maps in a Large Sample of Human Observers

**DOI:** 10.1101/2020.12.30.424856

**Authors:** Noah C. Benson, Jennifer M. D. Yoon, Dylan Forenzo, Stephen A. Engel, Kendrick N. Kay, Jonathan Winawer

## Abstract

How variable is the functionally-defined structure of early visual areas in human cortex and how much variability is shared between twins? Here we quantify individual differences in the best understood functionally-defined regions of cortex: V1, V2, V3. The Human Connectome Project 7T Retinotopy Dataset includes retinotopic measurements from 181 subjects, including many twins. We trained four “anatomists” to manually define V1-V3 using retinotopic features. These definitions were more accurate than automated anatomical templates and showed that surface areas for these maps varied more than three-fold across individuals. This three-fold variation was little changed when normalizing visual area size by the surface area of the entire cerebral cortex. In addition to varying in size, we find that visual areas vary in how they sample the visual field. Specifically, the cortical magnification function differed substantially among individuals, with the relative amount of cortex devoted to central vision varying by more than a factor of 2. To complement the variability analysis, we examined the similarity of visual area size and structure across twins. Whereas the twin sample sizes are too small to make precise heritability estimates (50 monozygotic pairs, 34 dizygotic pairs), they nonetheless reveal high correlations, consistent with strong effects of the combination of shared genes and environment on visual area size. Collectively, these results provide the most comprehensive account of individual variability in visual area structure to date, and provide a robust population benchmark against which new individuals and developmental and clinical populations can be compared.

**Significance Statement:** Areas V1, V2, and V3 are among the best studied functionally-defined regions in human cortex. Using the largest retinotopy dataset to date, we characterized the variability of these regions across individuals and the similarity between twin pairs. We find that the size of visual areas varies dramatically (up to 3.5x) across healthy young adults, far more than the variability of the cerebral cortex size as a whole. Much of this variability appears to arise from inherited factors, as we find very high correlations in visual area size between monozygotic twin-pairs, and lower but still substantial correlations between dizygotic twin pairs. These results provide the most comprehensive assessment of how functionally defined visual cortex varies across the population to date.

## Introduction

Maps are a canonical organizing principle of the cerebral cortex. Perceptual regions of the brain in particular are tiled with numerous areas each organized along specific dimensions. These maps include tonotopic maps (auditory cortex), somatosensory homunculi (somatosensory cortex), and retinotopic maps (visual cortex). The retinotopic maps of the visual system are particularly well-studied and serve as a model system for the study of cortical organization generally. In each hemisphere, the first three visual maps (V1, V2, and V3) are among the largest distinct regions of the human cortex, and each forms a complete representation of the contralateral visual hemifield (reviewed by Wandell and Winawer, 2011). These cortical representations distort the spatial layout of the visual field but largely maintain its topology, resulting in the phenomenon of cortical magnification (Talbot and Marshall, 1941; Daniel and Whitteridge, 1961), by which some parts of the visual field (*e.g.*, the fovea) are represented by substantially more of the cortical surface area per square degree than other parts (*e.g.*, the periphery).

Due to high interest both in the visual system and in properties of cortical maps, the retinotopic organization of the early visual cortex has been measured many times. These studies began with lesion patients and rough estimates of anatomy (Inouye, 1909; Holmes, 1918) and grew over time to include post-mortem studies of the stria of Gennari (Stensaas et al., 1974), lesion studies linked to anatomical MRI (Horton and Hoyt, 1991), PET studies (Fox et al., 1986), detailed but small-*n* studies of functional and anatomical MRI (Engel et al., 1994; Sereno et al., 1995; Engel et al., 1997), computational models of cortex from many observers (Schira et al., 2010; Benson et al., 2014; Wang et al., 2015; Benson and Winawer, 2018), and retinotopy studies with very large subject pools (Wang et al., 2015; Benson et al., 2018). Consistent among these studies is the observation that the topology of human retinotopic organization is highly conserved across subjects: the locations and boundaries of early visual areas correspond to consistent visual-field and cortical-anatomical features across subjects. However, substantial variation in the size and internal structure of the maps persists. **Figure 1** plots the outline of area V1 in the left hemispheres of two subjects from this study who exemplify this variation. The V1 of the subject on the right is approximately 3.5 times as large as that of the left subject (**Fig. 1A**), but both the organization of the maps and the overall (whole-hemisphere) cortical surface areas are similar in both subjects (**Figs. 1B**, **1C**). Both the causes and the consequences of V1’s large variability in size across the population are unknown. Although some previous evidence exists that variation in the sizes of early visual areas is related to behavior (Duncan and Boynton, 2003; Schwarzkopf et al., 2011), and to other structures in the visual pathways (Andrews et al., 1997), precise characterizations of these relationships are absent due to the large sample sizes they would require. Although some evidence of genetic control over the wiring of the visual system and of environmental control over the intra-cortical organization exist from studies of achiasmatic subjects (Hoffmann et al., 2012), subjects with albinism (Hoffmann et al., 2003), and case studies of subjects with atypical or damaged visual systems (Adams et al., 2007; Muckli et al., 2009), direct evidence that the sizes of visual areas are environmentally or genetically determined is scant due to the rarity of twin populations in visual neuroimaging research.

**Figure 1.**
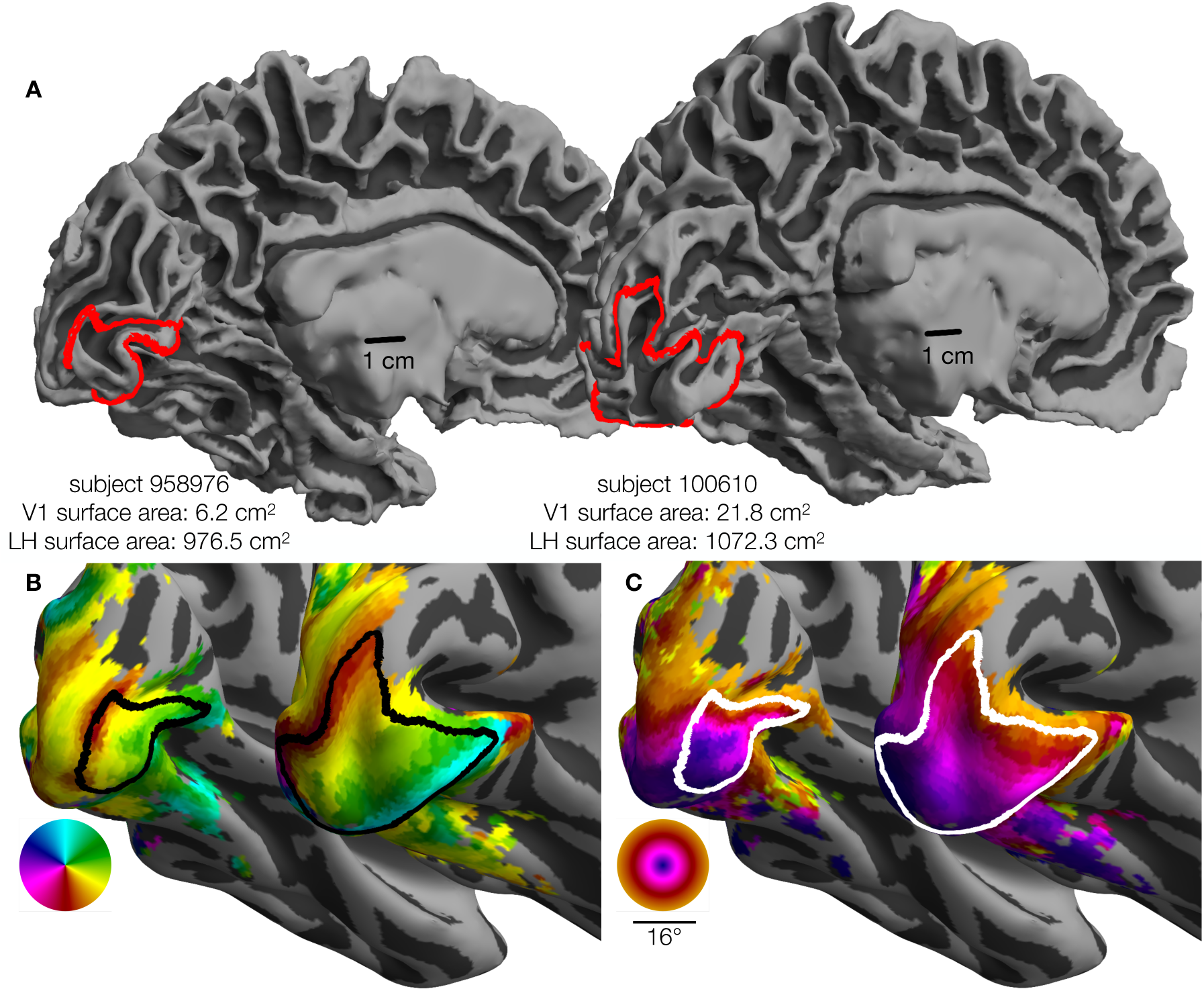
Variability in V1 size. The surface area of V1 (0-7°) differs by 3.5⨉ between two individuals. Each panel depicts the smallest and the largest LH V1 by surface area from the HCP 7T Retinotopy Dataset. In each panel, both hemispheres are rendered at the same scale. **A**. White matter surfaces, with the red line marking the V1 boundary (0-7° eccentricity). Identical 1 cm scale-bars are plotted at similar positions relative to each hemisphere. **B**. Inflated surfaces colored by polar angle; black lines mark the V1 boundary. **C**. Inflated surfaces colored by visual eccentricity; white lines mark the V1 boundary. Red, black, and white V1 boundaries are equivalent: Colors were chosen to improve contrast. Orthographic projections of these surfaces and boundaries that better show the context of the boundaries can be found in the project’s Open Science Framework repository in the “images/raw_lines” folder (see Methods).

Recently, the Human Connectome Project (HCP) (Van Essen et al., 2012) has published a huge trove of neuroimaging data, including retinotopic mapping data from 181 subjects (Benson et al., 2018). Details of these measurements and analyses have been described elsewhere (Van Essen et al., 2012; Benson et al., 2018), but, briefly: each subject participated in 6 distinct 5-minute population receptive field (PRF) scans at 7T, as well as detailed anatomical MRI. The retinotopic scans included expanding and contracting rings, clockwise and counterclockwise wedges, and sweeping bars with a maximum eccentricity of 8°. PRF models were fit for the whole brain. The 181 subjects included numerous monozygotic (MZ) as well as dizygotic (DZ) twin pairs, making it an unparalleled source of information about the variability of retinotopic organization. Here, we present our analysis of the variation in these maps across the subject population as well as comparison between MZ and DZ twin pairs.

To facilitate these comparisons, we manually labeled the V1, V2, and V3 boundaries, as well as the V1 horizontal meridian (HM) and five iso-eccentricity contours, in all 362 hemispheres. We then quantified the surface areas and cortical magnifications of the visual sectors and areas delineated by these contours. We also examined a variety of anatomical and functional properties such as the cortical gray-matter thickness and the population receptive field (PRF) parameters after aligning all subjects to a template surface in order to minimize anatomical variability across subjects. We have made this entire dataset of contours and surface areas publicly available with our analyses (https://osf.io/gqnp8/). Across our analyses, correlations of anatomical and functional measures between twin-pairs are high, and the similarity of MZ twins is significantly greater than that of DZ twins or unrelated pairs.

## Methods

All analyses in this paper were performed using the PRF analysis results of the HCP 7 Tesla Retinotopy Dataset (Benson et al., 2018). These PRF data include polar angle and eccentricity values for each vertex of each hemisphere of the 181 HCP subjects who participated in retinotopic mapping experiments. The analyses performed here consist of three steps: (1) drawing iso-angle and iso-eccentricity contours on each hemisphere, (2) pre-processing the contours and converting them from 2D image coordinates to 3D cortical surface coordinates, and (3) analyzing and comparing the visual regions defined by the contours. Of the 181 subjects in the dataset, 19 hemispheres (∼5%) were excluded due either to poor retinotopic map quality or to the inability of two or more “anatomists” to produce topologically valid sets of contours (see “Pre-processing the contours”, below).

### Experimental Design and Statistical Analyses

Experimental procedures for the HCP datasets have been described elsewhere (Glasser et al., 2016; Vu et al., 2017; Benson et al., 2018). Of the 181 HCP subjects, 109 were female and 72 were male. These include 53 pairs of genetically confirmed identical twins (106 individuals), 34 pairs of fraternal twins (68 individuals), two pairs of nontwin siblings (four individuals), and three individuals whose twins/siblings were not included. DZ pairs are all same sex.

### Drawing the Contours

The delineation of cortex was performed by four anatomists using custom MATLAB software. This software displayed an orthographic projection of a single spherical hemisphere at a time on which the anatomist could see various data displayed and could manually click points that define iso-eccentricity and iso-angle contours. We will refer to this projection of the cortical surface as the *anatomist projection*. The user was able to toggle between three data displays: the hemisphere’s polar angle map, eccentricity map, and binarized curvature map. The first two of these maps were obtained from the HCP 7T Retinotopy Dataset, while the curvature map comes from the HCP directly. Users could also toggle the display of contours delineating the anatomically-defined maximum probability map regions of the Wang *et al*. (2015) atlas. Using the software, anatomists could add and remove points that define a specific set of contours; these contours are described in **Table 1** and are shown in **Figure 2A**. Each anatomist indicated whether segments within any of their contours were low confidence (otherwise assumed high confidence), and additional notes could be added to the contours drawn for each hemisphere. We refer to one set of drawings of each of these contours on a single hemisphere as a “contour-set”.

**Figure 2.**
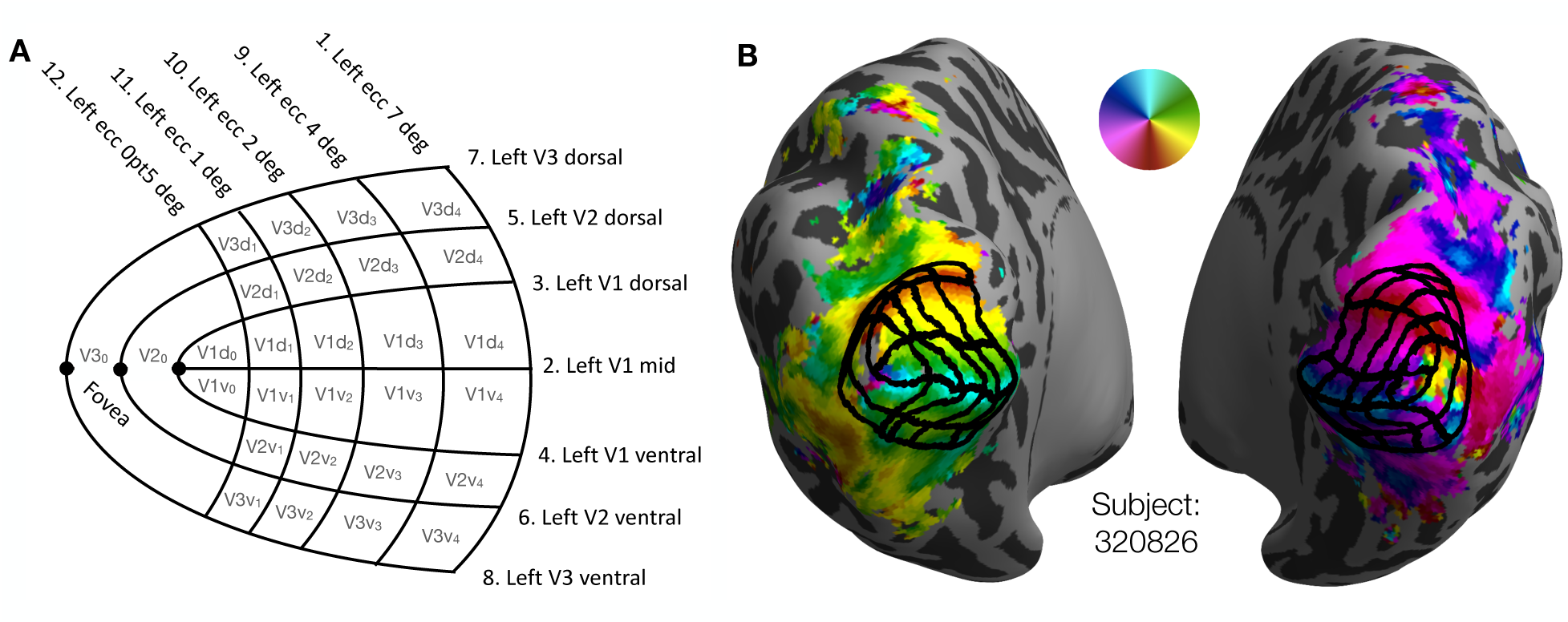
V1-V3 manually defined grids. **A**. Schematic of the manually defined V1-V3 grids marked by four human anatomists: 7 iso-polar angle contours (V1m, V1d, V1v, V2d, V2v, V3d, V3v) and 5 iso-eccentricity contours (0.5°, 1°, 2°, 4°, 7°) on each hemisphere (LH depicted). **B**. A subject’s inflated hemispheres with polar angle retinotopy (see color wheel), the manual definition of V1-V3 averaged across 4 anatomists (black grid).

**Table 1.** Contours drawn by anatomists on each hemisphere.

| Contour Name | Contour Description |
| --- | --- |
| V1 mid | The horizontal meridian in V1 |
| V1 dorsal | The lower vertical meridian, V1–V2 dorsal boundary |
| V1 ventral | The upper vertical meridian, V1–V2 ventral boundary |
| V2 dorsal | The lower vertical meridian, V2–V3 dorsal boundary |
| V2 ventral | The upper vertical meridian, V2–V3 ventral boundary |
| V3 dorsal | The lower vertical meridian, V3 dorsal–V3A/B boundary |
| V3 ventral | The upper vertical meridian, V3 ventral–hV4 boundary |
| 0.5° | The 0.5° iso-eccentricity contour through V1, V2, and V3 |
| 1° | The 1° iso-eccentricity contour through V1, V2, and V3 |
| 2° | The 2° iso-eccentricity contour through V1, V2, and V3 |
| 4° | The 4° iso-eccentricity contour through V1, V2, and V3 |
| 7° | The 7° iso-eccentricity contour through V1, V2, and V3 |

Anatomists were trained to draw the set of iso-angle and iso-eccentricity contours, shown as a schematic in **Figure 2A**, onto a subject’s polar angle and eccentricity maps. After each anatomist had drawn contours for a few subjects, their contours were reviewed by a senior author. Based on these reviews, we confirmed that all anatomists understood the task. The anatomists were then instructed to reproduce the set of contours on each subject’s hemisphere while preserving the topology of the contours found in **Figure 2A**—explicitly, each anatomist drew contours on each hemisphere. Anatomists were allowed to review and to make edits to previously drawn contours. Note that a core assumption of this method is that neurotypical adults will have retinotopic organizations in V1-V3 whose topology is compatible with that of the schematic. The job of the anatomists was to describe how the schematic mapped onto each hemisphere’s retinotopic maps, not to infer a novel topology. The extent to which this single topology is accurate for all subjects remains an open question.

### Pre-processing the Contours

The contours drawn by the anatomists were separated into distinct “contour-sets”: one set of 12 contours per anatomist per hemisphere (**Tab. 1**). Contour-sets were lightly edited by an automated algorithm designed to repair minor topological defects. These minor defects were limited to near-misses of iso-contours that should intersect—for example, if an anatomist were to have put the last point in the 4° iso-eccentricity contour just a few pixels away from the V3 dorsal contour at which it topologically terminates, then the algorithm would extend the contour to the appropriate terminus. Contour-sets that could not be repaired or that were discovered to be topologically incorrect—for example if the 2° and 4° iso-eccentricity contours intersected—were excluded from further analysis. These subjects and the reasons for their exclusion are included in the dataset on the OSF page (https://osf.io/gqnp8/).

After the contours were repaired, each contour was converted from a set of image coordinates to a “cortical path”. This conversion was performed by transforming the cortical surface mesh into the *anatomist projection* (see above, “Drawing the contours”), then converting each contour into a set of steps along the cortical surface by dividing each segment of the contour into smaller segments that each cross a single triangle of the mesh. The barycentric coordinates (*i.e.*, coordinates relative to the vertices of the triangle) of these smaller segments in the cortical surface mesh were then calculated and stored. The barycentric coordinates were used to project the complete contours onto other surface geometries such as the subjects’ white and pial surfaces as well as other registrations such as FreeSurfer’s *fsaverage* for comparison across subjects. Finally, each contour was divided into 499 segments of equal length with respect to the spherical *fsaverage* surface such that the 500 equally spaced points along its length could be averaged both across anatomists (for a single subject) or across subjects using the shared *fsaverage* registration. Average contours across subjects, across anatomists, and across both were calculated.

### Contour and Surface Area Analysis

The set of contours for each hemisphere parcellates the occipital region into three visual areas (V1, V2, V3) as well as several “sectors” within each of these areas. These sectors are labeled with gray text in **Figure 2A**, *e.g.*, “V3d1” for the dorsal portion of V3 within 0.5 and 1 deg eccentricity. Sectors were extracted from the contour sets for each anatomist by precisely dividing each triangle through which any contour passed into a set of sub-triangles inside the sector and a set of sub-triangles outside the sector. By doing this, exact surface areas of the sectors and visual areas drawn by the anatomists can be calculated rather than approximations based on vertex counts. This is important because triangles in the cortical meshes do not all have the same area.

Sectors for each hemisphere were projected onto the mid-gray cortical surface mesh for analysis of surface area and cortical magnification. Surface areas for each sector and visual area were calculated by summing the areas of the mesh triangles and partial sub-triangles they contained. No attempt was made to account for the curvature of individual triangles (*e.g.*, by using the vertex normals): surface areas were computed simply as the sum of areas of the triangles as they exist in the reconstructed surface. The cortical magnification of a sector was defined as its surface area divided by the area (in square degrees) of the visual field that it represents. This visual field area, for all sectors except the V1 and V3 foveal sectors (V20, and V30), is given by the formula *a* = *π* (*r*1^2^ - *r*0^2^) / 4 where *r*1 is the eccentricity of the outer contour and *r*0 is the eccentricity of the inner contour. For V20 and V30, which span both the upper and lower visual field, this equation must be multiplied by 2 and simplifies to *a* = (*π* / 8) deg^2^.

In addition to the cortical magnification of discrete sectors, a continuous version of cortical magnification *m*(*r*) [mm^2^/deg^2^] was calculated in terms of the eccentricity *r*. For a given visual area (V1, V2, or V3) and eccentricity *r*, *m*(*r*) is calculated by first finding the value Δ*r* such that 20% of the cortical surface vertices in V1, V2, or V3 have an eccentricity between *r* - Δ*r* and *r* + Δ*r*. The total cortical surface area represented by these vertices is then summed and divided by the visual area contained in the eccentricity ring defined by *r* ± Δ*r*.

We quantified the disagreement between anatomists at each point ***x*** along each of the 12 contours in the following way: First, for each subject, we computed the standard deviation *σ* of the positions of the equivalent points to ***x*** drawn by the four anatomists. Equivalent points are found by expressing points in terms of the fractional distance along their associated contours. Second, we averaged this quantity (*σ*) across the 181 subjects. This computation yields a single value at each point along each contour that is small anywhere anatomists agree and large anywhere anatomist contours diverge.

### Assessment of Heritability

Heritability was quantified first by comparing the correlations of the surface areas of V1-V3 of MZ twin-pairs to those of DZ twin-pairs. The intraclass correlation coefficient (ICC) was used (Shrout and Fleiss, 1979). The ICC of a set of paired values is similar to the Pearson correlation, but whereas the Pearson correlation measures the correlation of a set of *ordered* pairs, the ICC measures the correlation of a set of *unordered* pairs. Because it does not matter which member of a twin pair is “twin 1” and which member is “twin 2,” twins should be considered unordered pairs for the purposes of correlation. Specifically, we employ an unbiased estimate of the ICC that is derived from the one-way analysis of variance (McGraw and Wong, 1996). In this formulation, *r̂*_ICC_ = (MS*_r_* − MS*_w_*)/(MS*_r_* + MS*_w_*) where MS*r* and MS*w* are the mean-square of the across-pairs factor and the residuals, respectively. The formulae for MS*r* and MS*w* are given in **Eqs. 1** and **2**, respectively, where **X** is a 2×*n* matrix of twin-pair measurements such that *x*_1,j_ is the measurement of the first twin of twin-pair *j, x*_2,j_ is the measurement of the second twin, 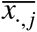 is the mean of *x*_1,_ *_j_* and *x*_2,_ *_j_*, and is the mean of 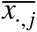 for *j* ∈ {1,2 . . . *n*.

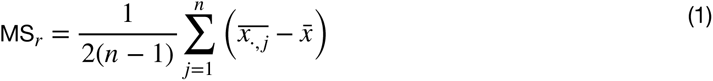

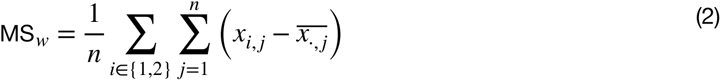

In principle, the computation of correlations in the size of a visual area between both MZ and DZ twins enables one to estimate the degree of heritability of that visual area’s size within the population, where heritability is the fraction of variance in the population for a given trait (*e.g.*, visual area size) that is partially attributable to genetics. In this study, we employed Falconer’s heritability index (*H^2^*), one of several formulas used in twin studies to estimate heritability (Falconer, 1960; Jacquard, 1983). Falconer’s formula is *H*^2^ = 2(*r*MZ - *r*DZ). Note that, with this formula, the confidence interval of the heritability will be larger than the confidence interval of the individual correlations (*r*MZ and *r*DZ) because variance is additive.

We additionally performed a comparison of the similarity in surface areas for MZ twin-pairs versus the similarity for DZ twin-pairs. We employed a non-parametric statistic for this evaluation because the absolute surface area differences between twin-pairs, being absolute, cannot be normally distributed. Although the differences may conform to the folded-normal distribution (Leone et al., 1961), they are incompatible with the multivariate ANOVA. For each MZ and each DZ twin-pair we calculated the surface area difference (in square cm) between the twins for the ROI combining V1-V3. We then asked whether, for a randomly selected MZ twin-pair, a randomly selected DZ twin-pair, and a randomly selected visual area (V1, V2, or V3), what is the probability that the MZ twin-pair has a smaller surface area difference with respect to the selected visual area than the DZ twin-pair? A probability value greater than 0.5 indicates an effect of twin type. The empirical probability of a bootstrap can be calculated by comparing every MZ twin-pair against every DZ twin-pair for each visual area. We computed the 2.5^th^, 50^th^, and 97.5^th^ percentiles of this probability using 10,000 bootstraps.

### Scientific Transparency and Software Availability

All data collected in this paper as well as all software tools used to analyze these data have been made freely available on an Open Science Framework website (https://osf.io/gqnp8/). Detailed instructions on how to access and analyze these data can be found there.

## Results

Within the 181 healthy, young adult subjects of the HCP 7T Retinotopy Dataset, the size of V1 varied dramatically. Defined from 0° to 7° of eccentricity, the smallest V1 was only 6.2 cm^2^, slightly more than a quarter the size of the largest V1 at 21.8 cm^2^ (**Fig. 1**). This 3.5-fold variation in the size of functionally-defined V1 is much greater than the variation in total hemisphere surface area across the same subjects, about 1.7-fold from smallest to largest. In the subsequent sections, we first evaluate the quality of the hand-drawn contours and use them to examine this variation. We then use the large number of twin pairs in the HCP 7T Retinotopy Dataset to quantify the similarity in the size of V1, V2, and V3 across monozygotic and dizygotic twin pairs.

### Hand-drawn Contours Are More Accurate than Anatomy-based Predictions and Show Agreement Across Anatomists

Delineation of visual area boundaries and iso-eccentricity contours is a difficult task. Across subjects, there is substantial variation in the anatomical arrangement of visual cortex and an approximately equivalent amount of variation in the functional maps themselves (Benson and Winawer, 2018). Beyond these variations, measurement noise and artifacts can cause large distortions in the maps. Although some work has been done to develop an automatic and objective method for drawing such contours (Dougherty et al., 2003; Benson and Winawer, 2018), these methods have been neither trained nor evaluated against a large gold-standard dataset such as the one described here. In light of these uncertainties, hand-labelling the V1- V3 contours is not only the most accurate available method of describing these ROIs, but it also provides a missing piece of any project to automate such description. Additionally, to mitigate sources of human error, we formalized a process by which four independent anatomists, using a common set of software, were trained and tested (see Methods).

Of the 1448 contour-sets, each of which comprises 7 iso-angle and 5 iso-eccentricity contours, 91 of them (∼6%) had some form of error and were discarded. Beyond these checks of basic correctness, we compared our hand-drawn contour-sets to the visual area boundaries obtained by anatomically-defined atlases that are based on FreeSurfer’s *fsaverage* alignment (Fischl et al., 1999), such as that of Wang *et al*. (2015). In some cases, the contours drawn by hand are very similar to those deduced by the atlas; however, when they differ, the hand-drawn contours match the functional landmarks of the visual area boundaries much more closely (**Fig. 3**). For example, in the top row of **Figure 3** (subject 221319), the dorsal V2 and V3 atlas boundaries deviate substantially from the functional polar angle reversals, while the hand-drawn contours match the reversals closely. In fact, for V3D, there is essentially no overlap between the hand-drawn and atlas-based maps.

**Figure 3.**
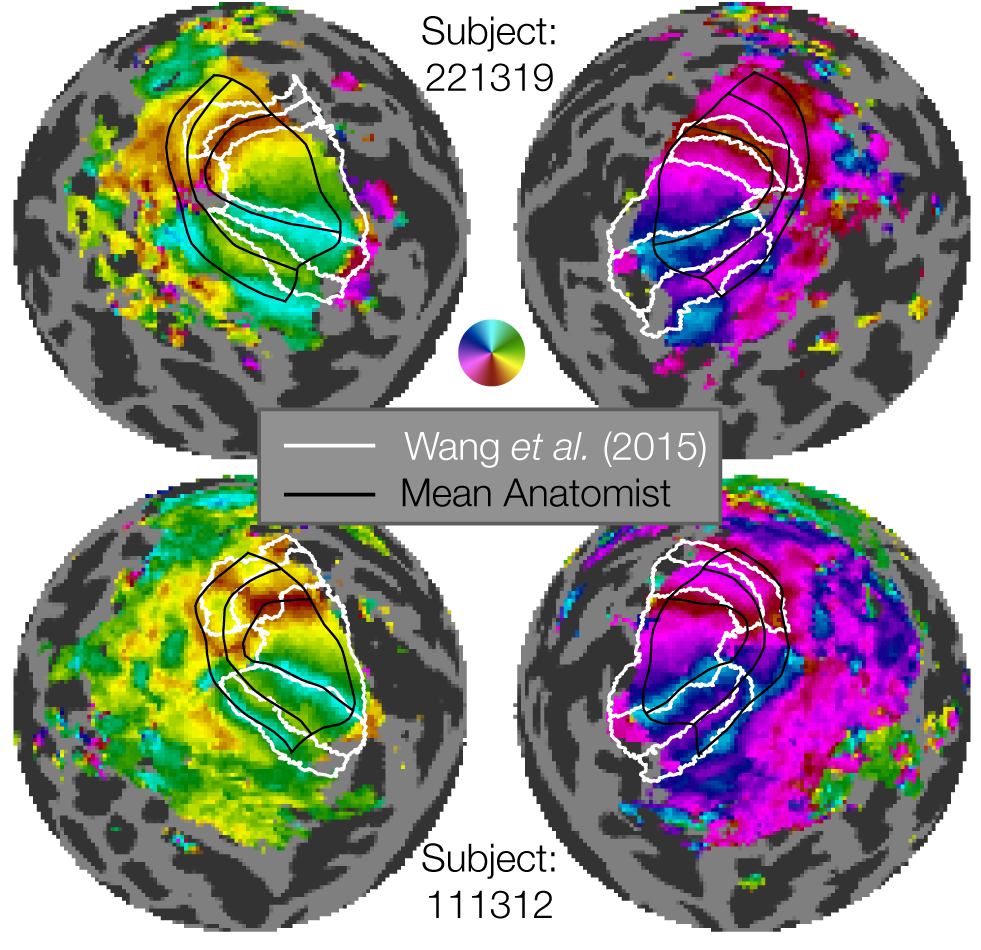
Examples of manually defined boundaries (black lines) compared to atlas-based boundaries (white lines). The figure shows flattened sphere representations of two example subjects. The top two hemispheres show an example of marked discrepancy between the manual definition and Wang atlas, especially in the dorsal regions of V1-V3. The bottom two hemispheres show a subject with substantial agreement between the manual definition and Wang atlas definition of V1v/d, V2v/d, V3v/d. The manually defined boundaries have the advantage of more closely capturing individual differences in retinotopy, as well as parcellating each area into multiple eccentricity sectors. Note that our manual definitions extend from 0-7° while the Wang atlas’s most foveal and eccentric boundaries differ (see text).

It is not surprising that the atlas-based contours will sometimes differ from landmarks in the functional data, as the atlas relies only on anatomy and the anatomical alignment method. These deviations have been quantified in prior work (Benson and Winawer, 2018). In comparison, the subject in the bottom row (subject 111312) has polar angle reversals that closely match the position of the visual area boundaries as predicted anatomically. (We do not compare the eccentricity contours since the atlas-based lines were fit to data spanning a different range of eccentricities than the hand-drawn contours.) Overall, we find that the hand-drawn contours procured by this project add substantial accuracy compared to anatomically- drawn contours when it comes to V1-V3 functional boundaries. These results underscore the importance of performing single-subject analyses that respect the features of individual subjects that may deviate from group atlases (such as surface-based *fsaverage* or volume-based MNI). The hand-drawn contours created here, although time-consuming and laborious, are critical for accurately measuring the structure of early visual areas, and this observation is likely to be the case for other areas in the brain. Similar renderings of atlas-drawn boundaries with hand-drawn contours for the 179 remaining subjects (both hemispheres, all anatomists) are available on the OSF site accompanying this publication.

An important test of the quality of the hand-drawn contours is the agreement across anatomists. First, we see that there is little systematic bias across anatomists. A comparison of each anatomist’s average contour across subjects reveals a high level of consistency (**Fig. 4A**). This consistency across anatomists is found even though there is substantial variation between subjects, evident in the plots of the average contour across anatomists per subject (**Fig. 4B**). Since these contours are all visualized after projection to the *fsaverage*, the spread among subjects indicates variability even after warping into this common space. Had we plotted the atlas-based contours such as those by Benson et al. (2014), Wang et al. (2015), and Glasser et al. (2016) in this space, they would be identical for each subject.

**Figure 4.**
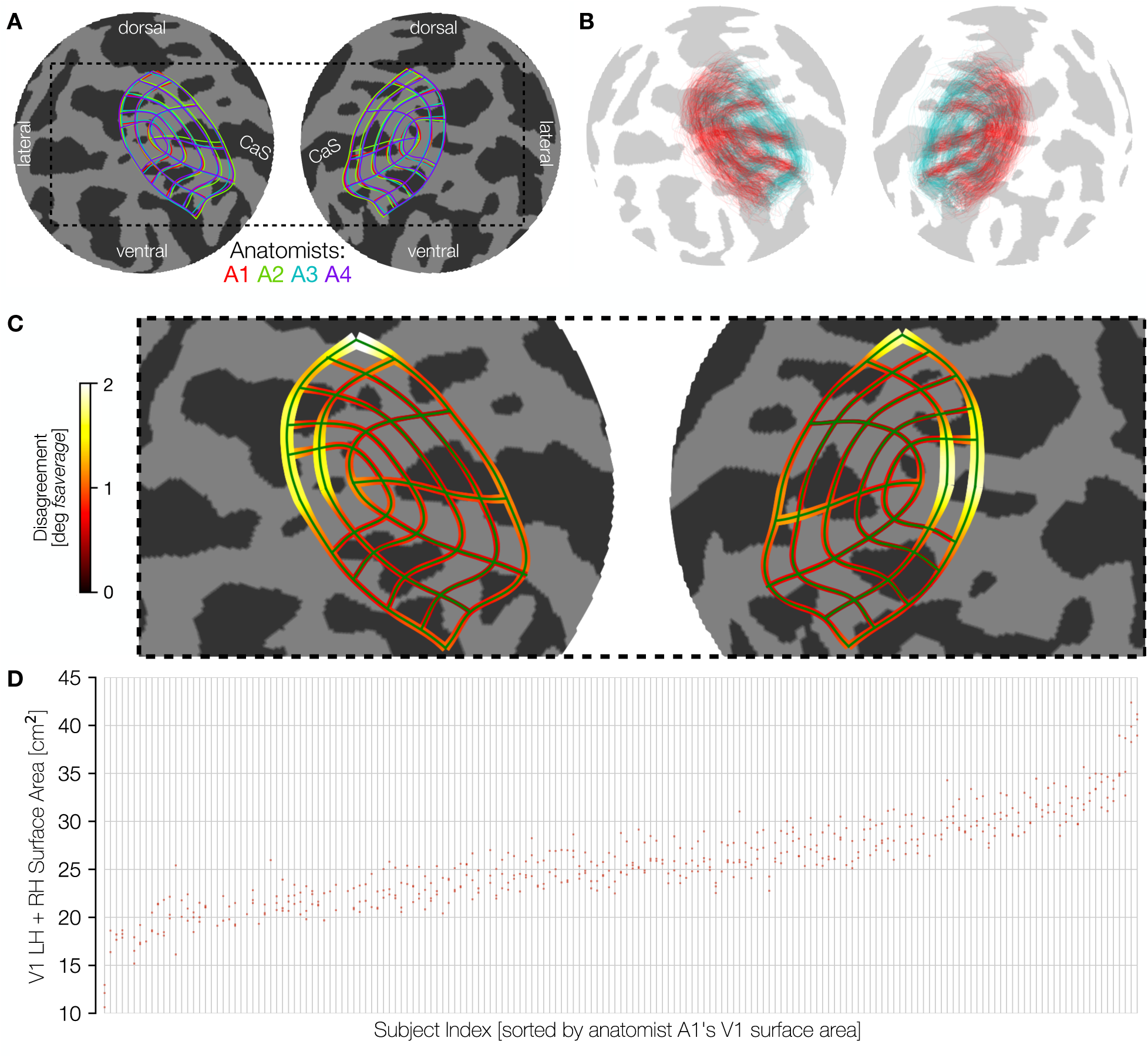
Consistency among anatomists and variability across subjects. **A**. The mean manually defined retinotopic grid (averaged across subjects) for each anatomist. Line thickness indicates ±*σ* across subjects for each anatomist. Plots use an orthographic projection of the *fsaverage* sphere centered at the occipital pole (CaS: Calcarine sulcus). **B**. Panel B is the converse of A: contours are averaged across the four anatomists, and plotted separately for each individual subject. For better visibility, we plot only a subset of the iso-eccentricity contours (0.5, 2, 7 degrees, colored cyan) and all 7 iso-polar angle contours (red). The variability across these retinotopic grid demonstrates inter-subject variability, even after warping to the *fsaverage* template. **C**. The magnitude of disagreement between anatomists is plotted at each point along an iso-eccentricity or iso-polar angle contour. Both the width of the contours and the color plot the mean over subjects of the disagreement between anatomists in degrees of the *fsaverage* spherical mesh. **D**. Surface area is plotted on the y-axis and individual subjects (light gray vertical lines) are ordered across the x-axis by sorting on A1 surface area. For each subject, the clustering is fairly tight for the four anatomists’ surface areas (red dots), and the spread between anatomists is much smaller than the magnitude of differences between subjects.

To quantify the consistency of the anatomists on a per-subject basis, we calculated the disagreement between anatomists at each point along each contour (see Methods). This disagreement is plotted in **Figure 4C**. Because disagreements were calculated in terms of distances along the surface of the spherical fsaverage mesh, these disagreements are in units of angular degrees of the sphere. These data demonstrate that the differences between anatomist contours are relatively low throughout most of early visual cortex with exceptions near the fovea and in dorsal V3. Even in these areas, the disagreement among anatomists averages only ∼2° of the spherical *fsaverage* cortical mesh.

The variability across subjects in the size of visual areas is much larger than the variability across anatomists. We show this by an intraclass correlation plot (**Fig. 3D)**. In this plot, subjects were sorted (along the *x*-axis) in ascending order by the bilateral surface area of the V1 ROI drawn by anatomist A1. The surface area measurements for all anatomists were then plotted vertically above the corresponding sorted *x*-position. In this plot the points are both tightly clustered in the *y*-dimensions and ascend smoothly across the *x*-dimension, indicating a high level of agreement between anatomists. The largest V1 (0 to 7°) is about 40 cm^2^ and the smallest about 12 cm^2^, whereas anatomist variability is only a few cm^2^ per subject. When the surface areas of V1 across all subjects and anatomists are fit with a general linear model in which each anatomist and each subject is represented by one factor, the factors associated with the subjects explain 93.9% of the variance. The remaining variance arises from factors associated with anatomists and the interaction between anatomists and subjects. In V2 and V3 these numbers are similar with 93.3% (V2) and 89.3% (V3) of the variance attributed to subjects. In short, the vast majority of the variance in our estimates of the size of all visual areas is due to subject variance, with variance associated with anatomists and subject- anatomist interactions both playing small roles.

### Cortical Surface Areas of V1-V3 Span a 3-Fold Range Across Subjects

The surface areas of V1-V3 vary far more between individuals than does total cortical area. Vertical histograms of the cortical surface area of V1, V2, and V3 across subjects are shown independently for each area, hemisphere, and sex in **Figure 5A**. These values are also documented in **Table 2**. See the OSF repository () for similar tables reporting visual area sizes derived from anatomically-defined boundaries (Benson et al., 2014) or extrapolation via cortical magnification. The largest to smallest V1 for males is about a 3.5 fold ratio, with a coefficient of variation (*σ* / *µ*) of 0.198. There are modest group differences in the distributions between males and females, and little to no differences between left and right hemispheres.

**Figure 5.**
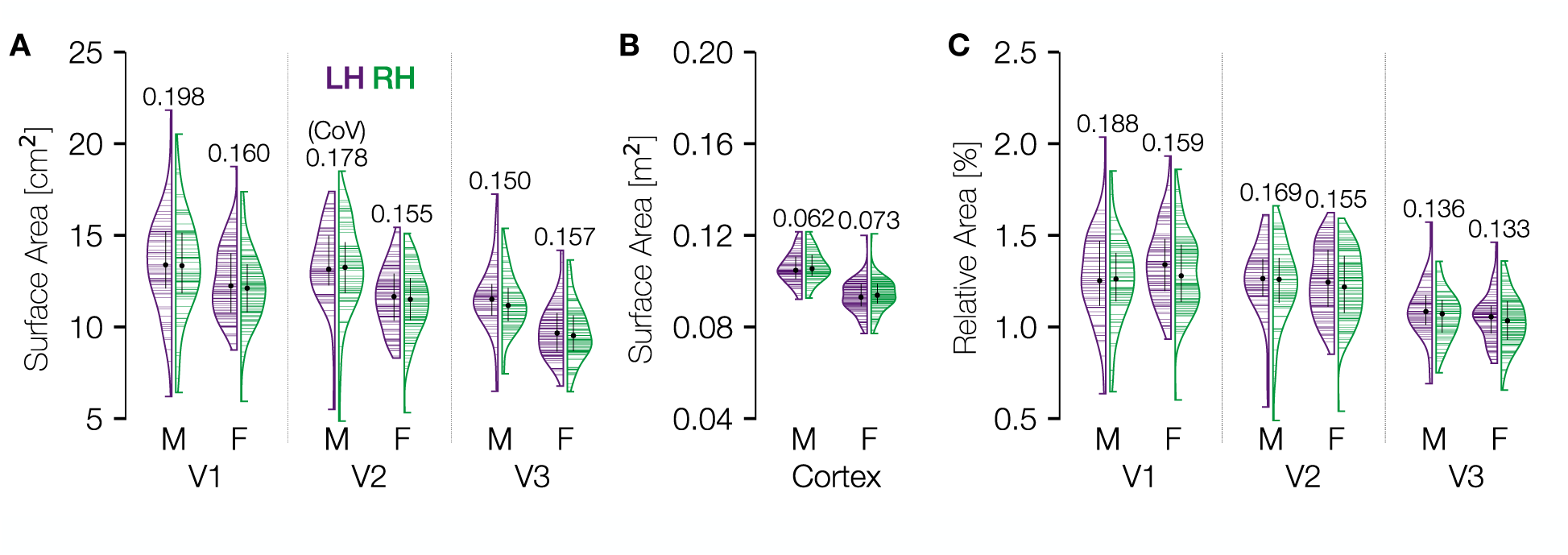
Distribution of cortical surface area in V1, V2, and V3. **A**. Violin plots of the total mid-gray cortical surface area (cm^2^) in areas V1-V3 for the 181 Human Connectome Project retinotopy subjects, defined from 0 to 7 deg eccentricity. Black dots and black lines embedded in the violin plots give the median ± interquartile range of the distribution while numbers above the plots give the coefficient of variation (standard deviation divided by mean). Horizontal lines indicate individual subjects. Distributions are plotted separateh by sex and laterality. **B**. Same as A but across the entire hemisphere. **C**. Same as A but normalized by total cortical area.

**Table 2.** Statistics of the surface areas of V1, V2, and V3.

| ROI | Sex | Hem. | Min | 25th% | Median | 75th% | Max | Mean | Std. Dev. | CoV | Max / Min |
| --- | --- | --- | --- | --- | --- | --- | --- | --- | --- | --- | --- |
| V1 | F | L | 8.7 (0.9%) | 10.8 (1.2%) | 12.2 (1.3%) | 14.0 (1.5%) | 18.8 (1.9%) | 12.5 (1.3%) | 1.9 (0.2%) | 0.16 (0.15) | 2.1 (2.1) |
|  |  | R | 6.0 (0.6%) | 10.8 (1.1%) | 12.1 (1.3%) | 13.4 (1.4%) | 17.4 (1.9%) | 12.2 (1.3%) | 2.0 (0.2%) | 0.16 (0.16) | 2.9 (3.1) |
|  |  | B | 17.1 (1.0%) | 22.2 (1.2%) | 24.7 (1.3%) | 26.9 (1.4%) | 36.1 (1.8%) | 24.9 (1.3%) | 3.5 (0.2%) | 0.14 (0.14) | 2.1 (1.9) |
|  | M | L | 6.2 (0.6%) | 12.1 (1.1%) | 13.4 (1.2%) | 15.2 (1.5%) | 21.8 (2.0%) | 13.5 (1.3%) | 2.7 (0.2%) | 0.20 (0.19) | 3.5 (3.2) |
|  |  | R | 6.4 (0.7%) | 11.8 (1.1%) | 13.3 (1.3%) | 15.1 (1.4%) | 20.5 (1.9%) | 13.4 (1.3%) | 2.6 (0.2%) | 0.20 (0.18) | 3.2 (2.9) |
|  |  | B | 12.6 (0.6%) | 24.3 (1.1%) | 27.2 (1.3%) | 30.4 (1.4%) | 40.4 (1.9%) | 27.1 (1.3%) | 5.1 (0.2%) | 0.19 (0.18) | 3.2 (2.9) |
|  | A | L | 6.2 (0.6%) | 11.2 (1.2%) | 12.8 (1.3%) | 14.4 (1.5%) | 21.8 (2.0%) | 12.9 (1.3%) | 2.3 (0.2%) | 0.18 (0.17) | 3.5 (3.2) |
|  |  | R | 6.0 (0.6%) | 11.1 (1.1%) | 12.5 (1.3%) | 13.9 (1.4%) | 20.5 (1.9%) | 12.7 (1.3%) | 2.4 (0.2%) | 0.19 (0.17) | 3.5 (3.1) |
|  |  | B | 12.6 (0.6%) | 22.8 (1.1%) | 25.6 (1.3%) | 28.6 (1.4%) | 40.4 (1.9%) | 25.8 (1.3%) | 4.4 (0.2%) | 0.17 (0.16) | 3.2 (2.9) |
| V2 | F | L | 8.3 (0.8%) | 10.4 (1.1%) | 11.7 (1.2%) | 12.9 (1.4%) | 15.4 (1.6%) | 11.6 (1.2%) | 1.8 (0.2%) | 0.15 (0.15) | 1.9 (1.9) |
|  |  | R | 5.3 (0.5%) | 10.4 (1.1%) | 11.5 (1.2%) | 12.7 (1.4%) | 15.1 (1.6%) | 11.5 (1.2%) | 1.8 (0.2%) | 0.16 (0.16) | 2.8 (3.0) |
|  |  | B | 17.0 (0.9%) | 20.7 (1.1%) | 23.4 (1.3%) | 25.8 (1.4%) | 30.3 (1.6%) | 23.3 (1.2%) | 3.2 (0.2%) | 0.14 (0.14) | 1.8 (1.7) |
|  | M | L | 5.5 (0.6%) | 12.3 (1.2%) | 13.2 (1.3%) | 15.0 (1.4%) | 17.4 (1.6%) | 13.3 (1.3%) | 2.3 (0.2%) | 0.17 (0.16) | 3.2 (2.9) |
|  |  | R | 4.9 (0.5%) | 11.9 (1.1%) | 13.2 (1.3%) | 14.6 (1.4%) | 18.5 (1.7%) | 13.2 (1.2%) | 2.4 (0.2%) | 0.18 (0.18) | 3.8 (3.4) |
|  |  | B | 10.4 (0.5%) | 24.3 (1.2%) | 27.0 (1.2%) | 29.9 (1.4%) | 35.2 (1.6%) | 26.6 (1.3%) | 4.5 (0.2%) | 0.17 (0.16) | 3.4 (3.0) |
|  | A | L | 5.5 (0.6%) | 10.9 (1.1%) | 12.3 (1.3%) | 13.7 (1.4%) | 17.4 (1.6%) | 12.3 (1.2%) | 2.1 (0.2%) | 0.17 (0.16) | 3.2 (2.9) |
|  |  | R | 4.9 (0.5%) | 10.7 (1.1%) | 12.1 (1.2%) | 13.7 (1.4%) | 18.5 (1.7%) | 12.2 (1.2%) | 2.3 (0.2%) | 0.19 (0.17) | 3.8 (3.4) |
|  |  | B | 10.4 (0.5%) | 22.0 (1.1%) | 24.6 (1.3%) | 27.3 (1.4%) | 35.2 (1.6%) | 24.7 (1.2%) | 4.2 (0.2%) | 0.17 (0.15) | 3.4 (3.0) |
| V3 | F | L | 6.8 (0.8%) | 8.7 (1.0%) | 9.7 (1.1%) | 10.8 (1.1%) | 14.2 (1.5%) | 9.8 (1.1%) | 1.5 (0.1%) | 0.15 (0.13) | 2.1 (1.8) |
|  |  | R | 6.5 (0.7%) | 8.7 (0.9%) | 9.5 (1.0%) | 10.6 (1.1%) | 13.7 (1.4%) | 9.7 (1.0%) | 1.5 (0.1%) | 0.16 (0.14) | 2.1 (2.1) |
|  |  | B | 14.3 (0.8%) | 17.6 (1.0%) | 19.5 (1.0%) | 21.2 (1.1%) | 27.6 (1.4%) | 19.6 (1.1%) | 2.9 (0.1%) | 0.15 (0.12) | 1.9 (1.8) |
|  | M | L | 6.5 (0.7%) | 10.6 (1.0%) | 11.5 (1.1%) | 12.3 (1.2%) | 17.2 (1.6%) | 11.6 (1.1%) | 1.9 (0.2%) | 0.16 (0.14) | 2.7 (2.3) |
|  |  | R | 7.5 (0.8%) | 10.3 (1.0%) | 11.2 (1.1%) | 12.1 (1.1%) | 15.4 (1.4%) | 11.3 (1.1%) | 1.5 (0.1%) | 0.13 (0.12) | 2.1 (1.8) |
|  |  | B | 14.2 (0.7%) | 21.2 (1.0%) | 22.9 (1.1%) | 24.6 (1.2%) | 32.6 (1.5%) | 22.9 (1.1%) | 3.2 (0.1%) | 0.14 (0.13) | 2.3 (2.0) |
|  | A | L | 6.5 (0.7%) | 9.2 (1.0%) | 10.5 (1.1%) | 11.7 (1.1%) | 17.2 (1.6%) | 10.6 (1.1%) | 1.9 (0.1%) | 0.18 (0.14) | 2.7 (2.3) |
|  |  | R | 6.5 (0.7%) | 9.1 (0.9%) | 10.3 (1.0%) | 11.5 (1.1%) | 15.4 (1.4%) | 10.3 (1.0%) | 1.7 (0.1%) | 0.17 (0.13) | 2.4 (2.1) |
|  |  | B | 14.2 (0.7%) | 18.5 (1.0%) | 20.8 (1.1%) | 22.9 (1.1%) | 32.6 (1.5%) | 21.0 (1.1%) | 3.4 (0.1%) | 0.16 (0.12) | 2.3 (2.0) |

“Sex” specifies females (F), males (M), or all participants (A). “Hemi.” specifies that the ROI is in the LH (L), the RH (R), or both (B). Columns 4-10 report cm<sup>2</sup>. Parentheses give results normalized by total cortical area. Similar tables based on the anatomically-defined boundaries or extrapolation to the full visual area using cortical magnification are provided in the OSF repository (DOI: 10.17605/OSF.IO/GQNP8).

Consistent with previous results (Dougherty et al., 2003), V2 is nearly as large as V1 (the median V2 surface area is 97% that of V1), and V3 is slightly smaller (17% smaller than V1). As a comparison we have also plotted the distribution of overall cortical surface area per hemisphere (**Fig. 5B**). Because the axes of **Figures 5A** and **5B** cover identical multiplicative ranges, it is readily apparent that the distribution of V1 sizes is much wider relative to its mean than that of the cortex overall. There is also a difference in overall cortical surface area between biological males and females. Normalizing each subject’s V1, V2, and V3 surface areas by their total cortical area removes the sex difference but has little effect on the variability per area within sex (**Fig. 5C).** Notably, the coefficients of variation (CoV) of the V1 distribution (∼0.2 and 0.16 for males and females) are not substantially affected by normalization (0.19 and 0.16 after normalization). There is also little change for V2 after normalization. In V3, the variability in surface area declines only slightly following normalization, suggesting that some of the variation in the size of V3 may be predicted by total cortical area.

### The Cortical Magnification Function Varies Substantially Across Subjects

The amount of cortical area devoted to different parts of the visual field varies systematically with eccentricity. Cortical magnification is the ratio of cortical area (e.g., in mm^2^) to visual area represented (*e.g.*, in deg^2^), and this value computed at each point in the visual field defines the cortical magnification function (CMF). This function is of considerable interest as a summary metric of map organization, and as a basis for comparison between individuals (Dougherty et al., 2003), between species (Horton and Hoyt, 1991), between different parts of the visual system (Adams and Horton, 2003), and between brain and behavior (Cowey and Rolls, 1974; Duncan and Boynton, 2003; Song et al., 2015).

Total cortical magnification can be calculated and reported in a number of ways. One highly robust way to calculate cortical magnification, given the many sectors into which we have divided the cortex, is to calculate the surface area of a particular sector and divide that area by the area of the region in the visual field that is represented by that particular sector. For example, in left V1, sector V1d2 (see **Fig. 2A**) is bordered in the visual field by the lower vertical meridian, the right horizontal meridian, the 1° iso-eccentricity arc and the 2° iso-eccentricity arc. The surface area of this partial annulus is ∼9.4 degrees^2^ (see Methods for more information).

The cortical magnification values for each sector, computed using this method, are reported in **Figure 6A**. These data confirm that the mean eccentricity follows the same basic p a t te r n a s h a s b e e n re p o r te d previously, with high magnification in the fovea gradually decreasing magnification in the periphery. In fact, the mean cortical magnification values in V1 are nearly identical to those measured almost 30 years ago by (Horton and Hoyt, 1991). To better visualize this, **Figure 6B** plots the distribution of cortical magnification across subjects in terms of eccentricity with the relationship reported by Horton and Hoyt plotted for comparison. In V1 in particular, the similarity of the median cortical magnification, as measured in 181 subjects using contemporary state-of-the-art methods, and the cortical magnification deduced 30 years ago through the examination of scotoma and lesions is remarkable. We can additionally observe the crossover in cortical magnification between V1 and V2/V3 between 0.5° and 1° of eccentricity previously observed by (Schira et al., 2010). At eccentricities higher than this crossover point, V1 has a higher cortical magnification than V2 and V3 while at eccentricities lower than the crossover V1 has the lower magnification.

**Figure 6.**
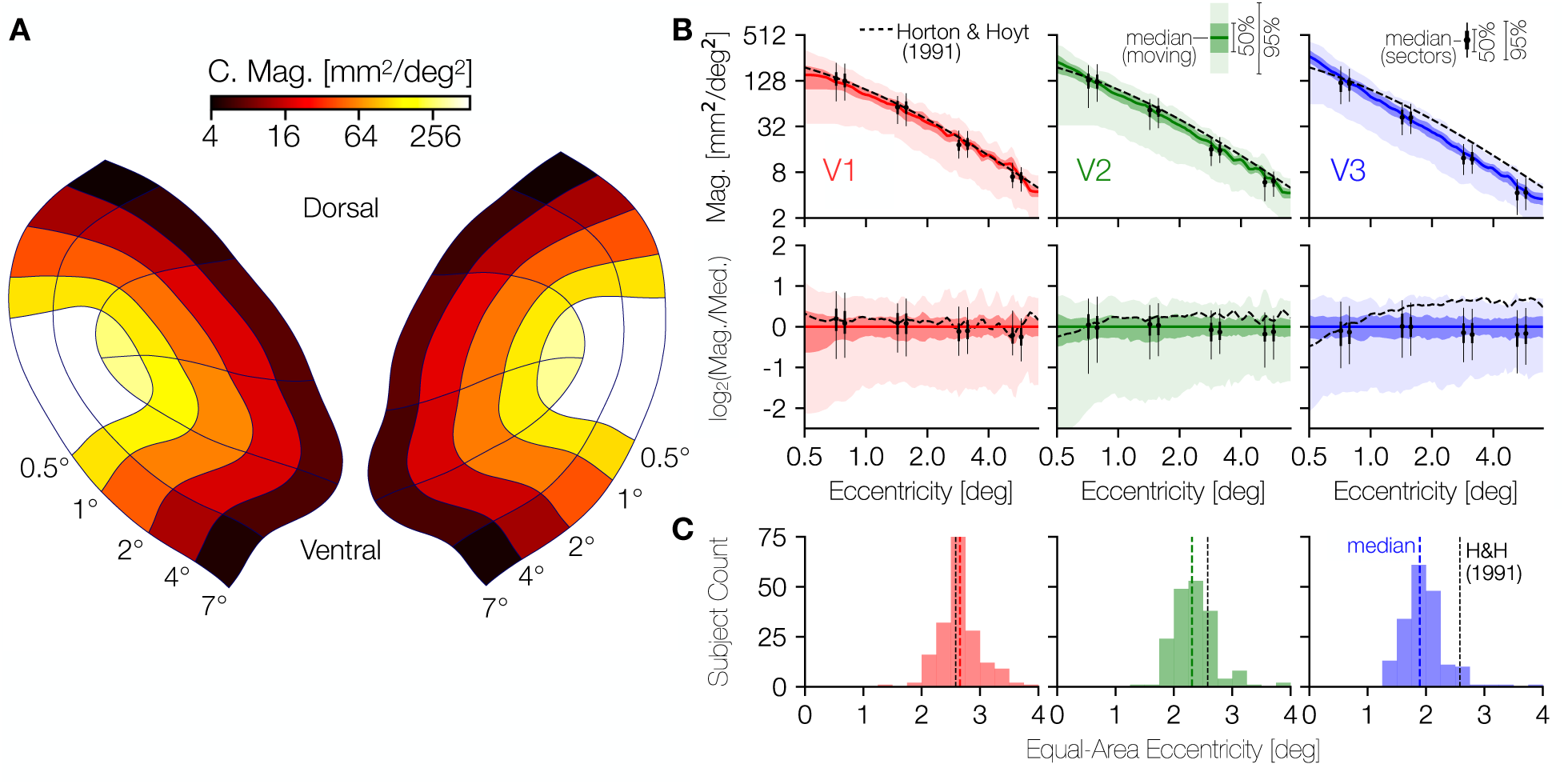
Cortical magnification in V1, V2, and V3. **A**. Average areal cortical magnification of each sector of V1-V3 across subjects; the mean set of contours across anatomists for each subject was used to calculate the cortical surface area of each sector. **B**. Median areal cortical magnification across subjects in terms of eccentricity. In both top and bottom rows, solid colored lines indicate the median while the darker and lighter shaded regions depict the 25%/ 75% and 2.5%/97.5% percentiles. The black dashed line represents the cortical magnification function for V1 reported by Horton and Hoyt (1991). In the top row of panels the cortical magnification is plotted. In the bottom row, the same data are reproduced but are plotted in terms of their relation to the median. A point (*x*_top_,*y*_top_) from the top row is plotted in the bottom row as (*x*_bot_,*y*_bot_) where *x*_bot_ = *x*_top_, but with *y*_bot_ = log_2_(*y* / *m*(*x*)) where *m*(*x*) is the median cortical magnification at eccentricity *x*. A consequence of this scaling is that the median line in the top row is plotted as a constant *y* = 0 in the bottom row. **C**. Histograms of the eccentricity that divides the surface area of the inner 7° of V1, V2, or V3 into equal foveal and peripheral halves. The black dotted line indicates the eccentricity at which this would occur according to Horton and Hoyt (1991): ∼2.58° of eccentricity. Explicitly, this means that, according to Horton and Hoyt, the V1 ROI from 0°-2.58° of eccentricity has the same surface area as the V1 ROI from 2.58°-7° of eccentricity.

Interestingly, the variability of cortical magnification is much higher near the fovea (<1° of eccentricity) in V1 and V2 than in the periphery, as can be seen most clearly in the second row of **Figure 6B**. This row plots the identical cortical magnification data as the top row, but plots it as a ratio to the median cortical magnification. *I.e.*, the *y*-axis is in log2 steps, so a *y*- value of 1 indicates twice the value of the median while a *y*-value of -1 indicates half the value of the median.

Notably, across visual areas and eccentricities, the variability in cortical magnification is commensurate with or slightly larger than the 3.5-fold variability across visual area surface areas. Some of the variability across subjects in cortical magnification values is due to variation in the overall size of V1 across subjects; other variability is likely related to the shape of each subject’s cortical magnification function. For example, two subjects with identically sized V1 regions may nonetheless devote very different fractions of their V1s to the inner 5° of eccentricity. In **Figure 6C** we report a model-free metric of the “steepness” of the cortical magnification that is independent of total visual area size: the eccentricity that splits the ROI (*i.e.*, the part of V1, V2, or V3 traced by the anatomists) into two ROIs with equal surface area. The eccentricity predicted by the measurements of Horton and Hoyt for this same ROI in V1 is also included as a vertical dashed line in each panel (∼2.58°). As one moves from V1 to V2 to V3, the median halfway-point eccentricity decreases by ∼1°. The halfway-point varies by a factor of approximately 2 in each visual area and is significantly correlated across hemispheres for each visual area (*r*V1 = 0.44, 95% CIV1 = 0.29–0.56; *r*V2 = 0.39, 95% C1V2 = 0.23–0.53; *r*V3 = 0.45, 95% C1V3 = 0.31–0.56; using 10,000 bootstraps). This indicates that the variation we report across individuals is unlikely to simply be due to measurement noise.

### Surface Area is Correlated Between ROIs, Hemispheres, and Dorsal/Ventral Subdivisions

Models of V1-V3 frequently include the assumptions that these visual areas are, to a first approximation, left-right and ventral-dorsal symmetric and that V1, V2, and V3 are closely related in size. Across subjects, we find this latter statement to be largely true. The surface area of V1, summed across hemispheres, is highly correlated with the surface area of V2 (*r* = 0.77, 95% C.I. = 0.73-0.81 across 10,000 bootstraps) (**Fig. 7A, 7D**). There is also a strong correlation between the surface areas of V2 and V3 (*r* = 0.64), and a lower but still robust correlation between V1 and V3 (*r* = 0.45). These correlations are higher than those reported previously (Dougherty et al., 2003), particularly the relation between V1 and V3 surface area which has been reported to be near 0. One reason we report larger correlations is in the methodology: we summed across hemisphere and across ventral and dorsal subdivisions, whereas Dougherty *et al*. did not. This summation may attenuate the effect of measurement noise. Secondly, the much larger sample size available from the HCP data affords a more accurate estimate of the true population relationships.

**Figure 7.**
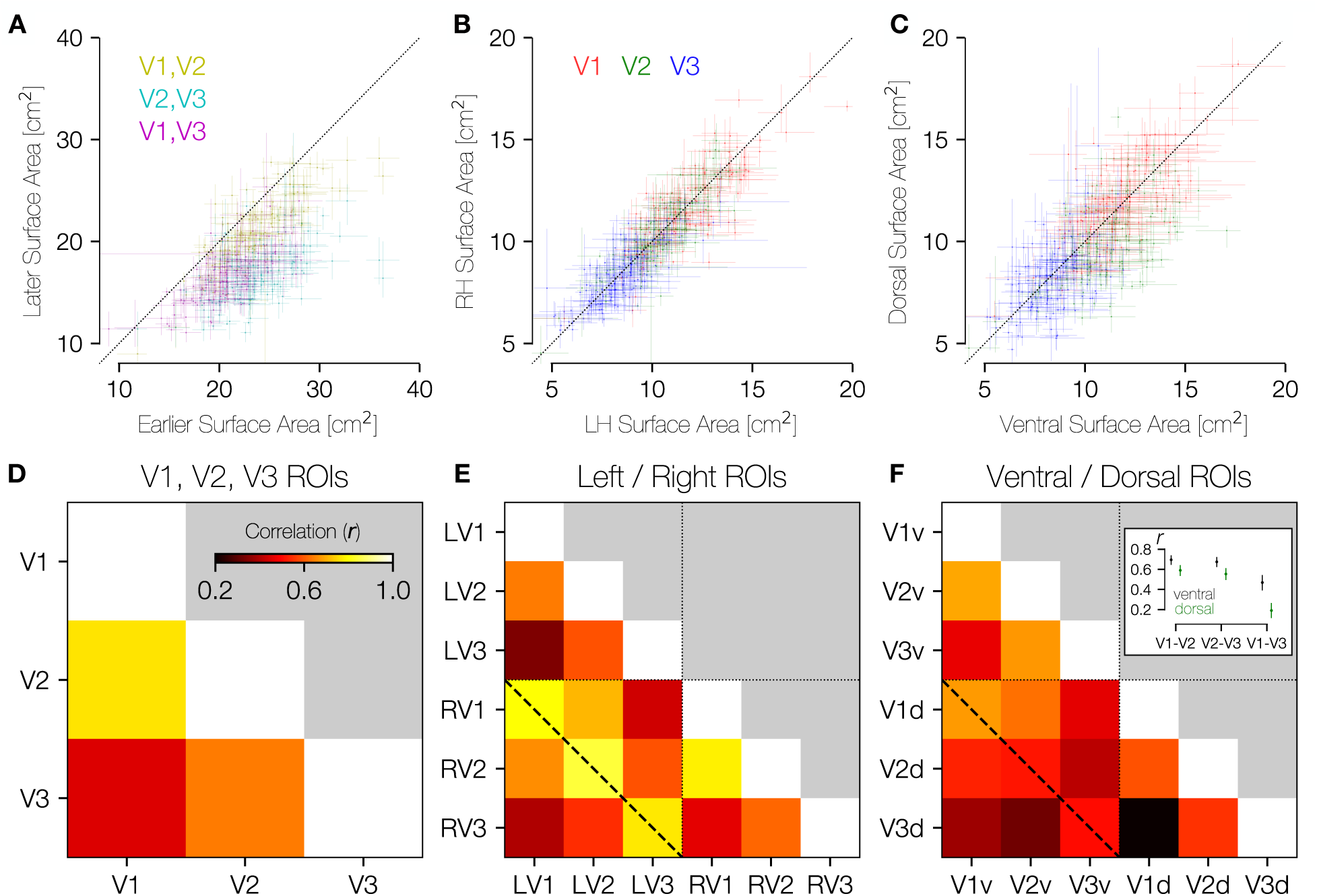
Area-area correlations. **A**. In all three possible pairings of areas V1, V2, and V3, the earlier area tends to be larger than the later area (center of mass for all three colors lies below the lines of equality). Crosses about the plotted points indicate the range of values across anatomists **B.** Within individuals, there is substantial LH-RH symmetry in the surface area of V1-V3. **C**. Within individuals, there is substantial dorsal-ventral symmetry in surface area. **D**. The correlation between areas V1 and V2 is higher than between areas V1 and V3 or areas V2 and V3. **E.** This correlation matrix includes LH/RH V1, V2, V3. Relatively higher correlation in surface areas can be seen within each area across hemispheres (e.g., LH V1 and RH V1; diagonal dotted line). **F**. This correlation matrix includes ventral/ dorsal V1, V2, V3. The pattern of correlations in surface area is highest for V1v-V2v, V2v-V3v, and V1v-V1d. The inset panel shows the median and 68% confidence intervals for ventral and dorsal comparisons, as computed across 10,000 bootstraps. Confidence intervals are non-overlapping for each comparison shown.

Within each of V1, V2, and V3 the surface areas of the right and left hemispheres are highly similar. **Figure 7B** shows a scatter-plot of the LH versus RH surface areas of V1, V2, and V3 for all subjects, which reveals the strong correlations as well as the overall decrease in size from V1 to V2 to V3. A similar plot is shown in **Figure 7C**, which compares the ventral surface area of V1-V3 (*x*-axis) to the dorsal surface area (*y*-axis). Again, there is high correlation within areas, and one can observe the decrease in size from V1 to V3. Unlike left versus right, these dorsal-ventral plots show some asymmetries: The points tend to be under the diagonal in V2 (*µ*V2v = 11.23 cm^2^ > *µ*V2d = 9.66 cm^2^), with V1 and V3 being more symmetrically arranged (*µ*V1d = 11.73 cm^2^ ≈ *µ*V1v = 11.37 cm^2^; *µ*V3d = 8.25 cm^2^ ≈ *µ*V3v = 8.34 cm^2^). (At a finer angular resolution than quarterfields, there are additional polar angle asymmetries in V1) (Benson et al., 2020). The differences between visual areas may reflect different processing priorities, and warrants further study.

Correlations across subjects for left v. right visual areas are strongest for corresponding areas. In **Figure 7E** we plot the correlation matrix that results from comparing LH size to RH size across subjects. The dashed line in this plot that marks the correlations between the same areas in contralateral hemispheres shows the highest correlations (all > 0.77). In comparison, the correlations shown between ventral and dorsal subparts of each visual area (**Fig. 7F**) show much lower (though still positive and highly significant) correlations. In fact, the highest correlations in this comparison are between V1-ventral and V2-ventral and between V2-ventral and V3-ventral as well as between V1-ventral and V1-dorsal. Correlations between equivalent dorsal regions are much lower (**Fig. 7F**, inset), suggesting either that processing needs for the ventral portions of V1-V3 (upper visual field) could be more similar across areas than those of the dorsal portion (lower visual field) or that dorsal areas are subject to greater measurement noise.

### The Size of Retinotopic Maps is Similar Between Twin Pairs

Datasets containing twin pairs are rare in visual neuroscience; accordingly, a uniquely valuable component of the dataset described in this paper is the availability of both MZ and DZ twin pairs (50 MZ pairs and 34 DZ pairs). We exploit these twin pairs to examine the extent to which variability in visual area size is explainable by relatedness. If twins have highly correlated surface areas, and the correlations are higher for MZ twins than DZ twins, it suggests that to a large degree, the size of these visual areas is inherited. Correlations that are high for both types of twin pairs but show little difference between the two types implicate a large role for shared environment in determining visual area size.

Visual inspection of the contours drawn for pairs of twins and pairs of unrelated subjects suggests that twins tend to have more similar map organization than unrelated pairs (**Fig. 8**). Although the flattened cortical maps introduce a small amount of arbitrary distortion in each hemisphere and thus should not be interpreted as a perfect proxy for similarity, it remains clear from examining both the contour geometry and the map parameters that each subject’s retinotopic organization is more similar to that of their associated twin than to that of either of the unrelated subjects. For example, the MZ twins in **Figure 8** both show relatively large foveal sectors in V2 and V3, differing from the two DZ twins.

**Figure 8.**
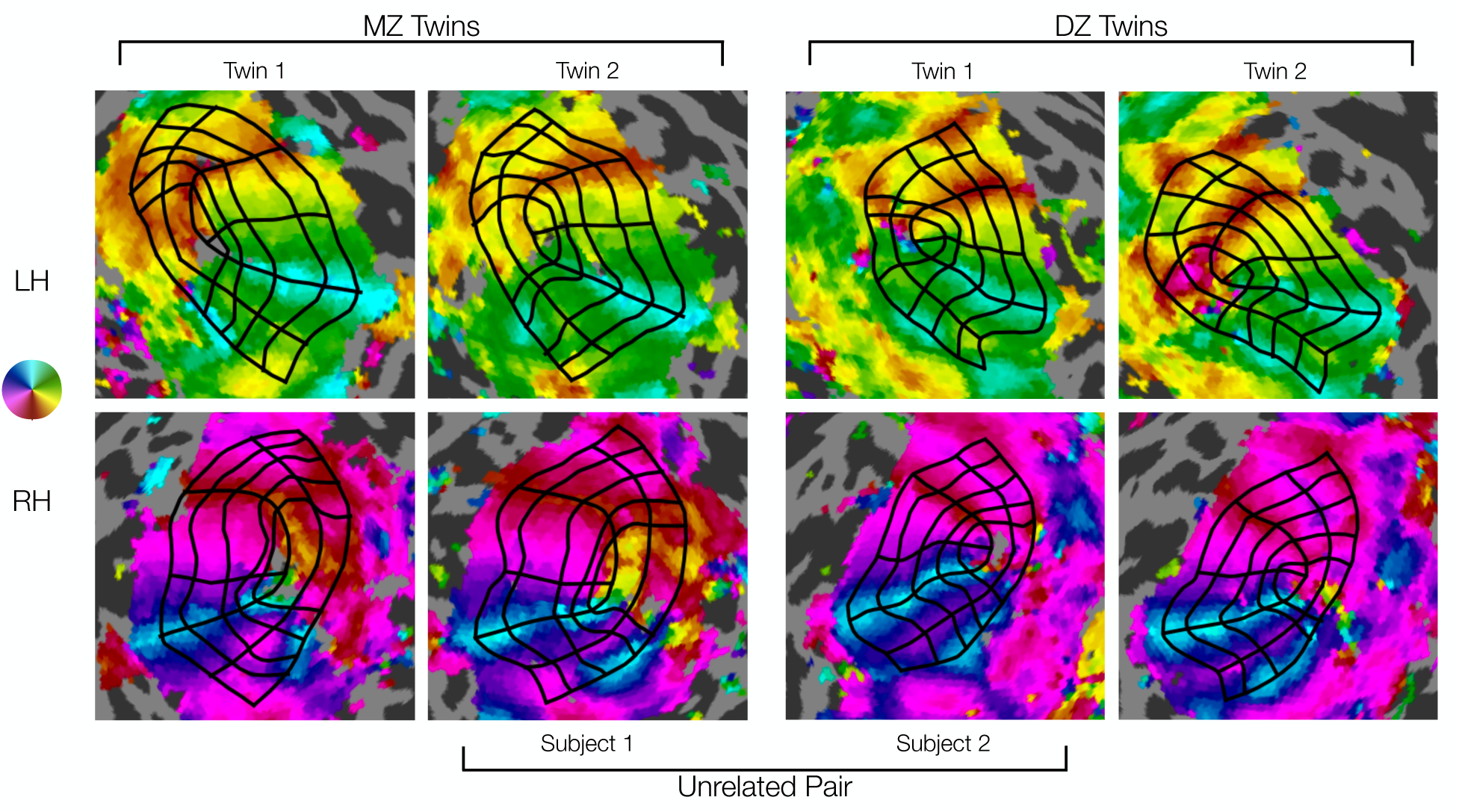
Comparison of manually defined retinotopic grids in MZ twins, DZ twins, and an unrelated pair. The top row shows left hemispheres and the bottom row shows right hemispheres. Functional data from polar angle scans is depicted for each hemisphere (see colormap). Black grids indicate the 7 iso-polar angle contours (V1m, V1d, V1v, V2d, V2v, V3d, V3v) and 5 iso-eccentricity contours (0.5, 1, 2, 4, 7 degrees) on each hemisphere (see Figure 2A). Note the high similarity in the functional data and resulting grid in the MZ twins (left two columns), especially the RH V1’s larger lower field (dorsal) representations. Intermediate similarity in the DZ twins (right two columns) can be seen in size, shape, and other details of the functional data and resulting grid. Lower similarity in the example unrelated pair (inner two columns) can be seen in the functional data and resulting grids, especially in overall size. Visual inspection reveals much higher similarity in the topography of areas V1-V3 for the DZ and MZ twins.

To quantify the similarity between sets of twin-pairs, we first computed the intraclass correlation coefficient (ICC) of the surface areas. We focus on the surface areas of V1, V2, and V3 because surface area is a well-defined proxy for the overall organization of these visual areas that can be derived from the drawn contours and because we have already established that surface area has a large amount of individual variability that is not due to measurement noise.

All three visual areas show very high correlations between MZ twins, with *r*ICC of 0.84, 0.81, and 0.75 for V1, V2, and V3, respectively (**Figs. 9A**, **9B**, and **9C**). This indicates that twin pairings explain about 80% of the variance in the size of these visual areas within the group of all MZ twins. We find lower but highly robust correlations for the DZ twins, with *r*ICC of 0.68, 0.72, and 0.44 for V1, V2, and V3. These calculations are based on the normalized surface areas (surface area of visual area divided by surface area of all of cerebral cortex); similar values are obtained for the unnormalized surface areas, with the only substantial exception being that the *r*ICC for V3 in the DZ twins is higher for the unnormalized calculations (0.72 v. 0.44). This indicates that with the exception of V3 in the DZ twins, the correlations between twins are unrelated to the similarity in total cortical area.

**Figure 9.**
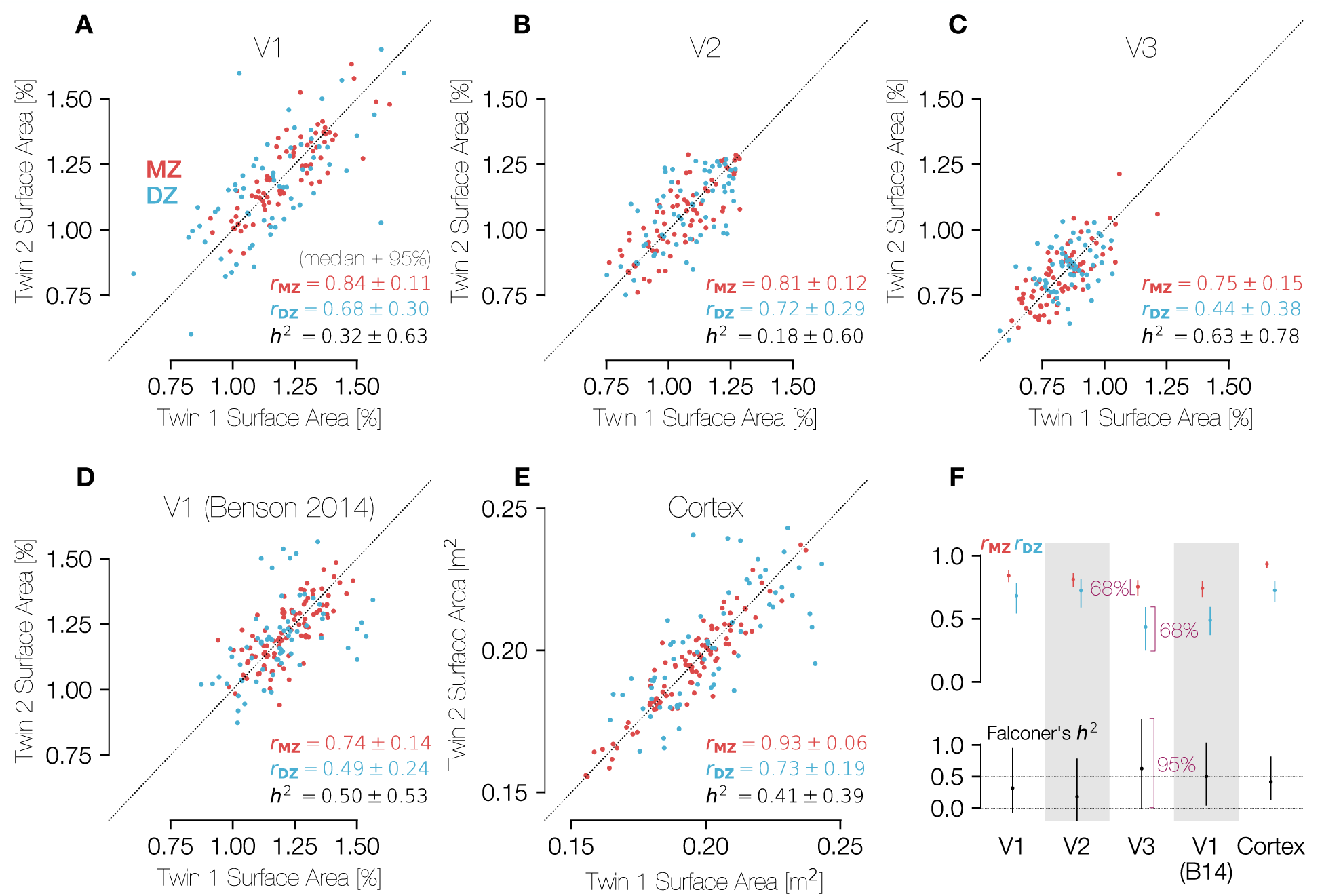
Correlations in the sizes of brain areas across MZ and DZ twins. The three panels in the top row show comparisons of the (**A**) V1, (**B**) V2, and (**C**) V3 percent-of-cortical surface area measurements for both MZ and DZ twin pairs. Because the assignment of a subject into the category of “twin 1” or “twin 2” is entirely arbitrary, all points plotted twice, reflected along the line x = y. Panel D plots the same metric for the atlas V1 boundaries of Benson et al. (2014), which are obtained through anatomical alignment and thus represent an anatomical definition of V1. In panels A-D, the surface areas were calculated between 0.5° and 7° of eccentricity only. Panel **E** shows a similar comparison of the entire cortical surface area across twin pairs in terms of square meters. **F**. Summary of correlations and heritability calculations for each ROI and relationship type (MZ/DZ). The *H*^2^ values were calculated using Falconer’s formula of heritability. Confidence intervals (CIs) were calculated using 1000 bootstraps over twin pairs. For the top row, CIs are at the 68% level (1 S.E.M.), and for the bottom row they are at the 95% level. In panels A-E, CIs are expressed as “± *x*” where *x* is half of the difference between the 97.5th and the 2.5th percentiles.

Although the similarity between twin pairs in the sizes of V1, V2, and V3 is not explained by similarity in total cortical area, the total cortical area is in fact highly similar between twin pairs. Notably, the correlation for the full cortical surface is substantially higher than that for any other region and is substantially higher in MZ twins (*r*ICC = 0.93) than in DZ twins (*r*ICC = 0.73), consistent with prior reports showing high heritability of cortical surface area (Panizzon et al., 2009; Winkler et al., 2010; Gomez-Robles et al., 2015; Schmitt et al., 2019) (**Fig. 9E**).

We used the correlation values for each visual area and twin type to calculate Falconer’s heritability index *H*^2^ for each area (see Methods). The fact that the correlations between twin pairs was positive for both twin types and higher for the MZ twins than the DZ twins means that the *H^2^* value is positive for all metrics (V1, V2, and V3 surface areas, as well total cortical surface area; **Fig. 9F**). Nonetheless, it is clear from the large size of the 95% confidence intervals that the number of twin pairs is far too low to make a reliable estimate of *H^2^*. For example, our 95% confidence intervals for V1 span 0 and 100%. This contrasts with the measures of correlation *r*ICC, which have tighter confidence intervals. Specifically, the variance in *H^2^* is four times the sum of the two variances in the *r*ICC calculations.

As an alternative to directly calculating the heritability of V1-V3 surface areas using correlations, we can instead assess the median difference in surface areas for MZ twin-pairs (Δ̃_MZ_) compared to the median difference for DZ twin-pairs (Δ̃_DZ_) and the probability that Δ̃_MZ_ < Δ̃_DZ_. This calculation is a non-parametric homolog to a two-way ANOVA, which cannot be used because the surface area difference values are absolute and thus not normally distributed (see Methods for details). Across 10,000 bootstraps, we found that the surface area of V1-V3 differs by 4.33 cm^2^ (median across bootstraps) for MZ twin pairs, and 6.06 cm^2^ for DZ pairs. This was a highly reliable difference: the 2.5^th^ and 97.5^th^ percentiles of the differences between MZ twin pairs were 3.72 cm^2^ and 4.96 cm^2^, and for DZ twins 5.25 cm^2^ and 7.00 cm^2^. Overall, this means that the visual area size differed between DZ twins by about 41% more than it differed between MZ twins (95% CI of 22% to 72%). For both types of twin pairs, the visual area sizes were much more similar than they are for unrelated subject pairs, whose V1- V3 size differs by 10.10 cm^2^ (2.5^th^ and 97.5^th^ percentiles of 9.94 and 10.26 cm^2^). For these values, V1, V2, and V3 surface areas were summed into a single ROI that was not normalized by total cortical surface area. However, the effect of DZ differences < MZ differences < unrelated pair differences remains significant even when V1-V3 size is normalized by cortical size.

For comparison with the functionally-defined regions (V1, V2, and V3), we also computed the correlation across subjects of the surface area of a similar anatomically-defined region, the V1 ROI as originally defined by Benson *et al*. (2014) and updated by Benson and Winawer (2018) (**Fig. 9D**). Because this ROI is computed using purely anatomical data from the subjects, the correlation of its size between twins reflects only the similarity of the cortical folding pattern in the V1 region. The “Benson 2014” V1 ROI was limited to have the same eccentricity boundaries (0.5°-7°) as the hand-labeled ROIs compared in panels A-C. These anatomically-defined regions were found using FreeSurfer’s anatomical alignment and were not informed by functional data (*i.e.*, retinotopy measurements of these subjects). The correlations for the anatomically defined V1 are lower than those for functionally defined region (0.75 v. 0.84 and 0.49 v. 0.68: anatomically v. functionally-defined V1 for MZ and DZ twins respectively).

Moreover, if we compute the residual surface area, meaning the area of the functionally defined V1 minus the area of the anatomically defined V1, we find that these residual areas are also highly correlated between twins (0.64 and 0.39, MZ, and DZ). This indicates that the hand-drawn contours capture meaningful features of the maps. If the hand-drawn contours differed from the anatomical boundaries only due to noise, then the residuals would not be correlated between twin pairs.

### The Organization of V1-V3 is Similar for Twin Pairs and More Similar for MZ Pairs

The size of a visual area on cortex is one measurement of early visual cortical organization; however, the visual area size provides relatively little information about similarities in the relationship between anatomy and function or in the idiosyncrasies of the layout of the retinotopic maps on cortex. To better understand what aspects of the structure-function relationship of visual cortex are shared among twins, we also examined the correlation of various anatomical and functional properties within the occipital cortex between MZ twin-pairs, DZ twin-pairs, or UR (unrelated) pairs. In the calculations of correlation of an ROI’s surface area between twins, as performed in the previous section (**Fig. 9**), there is one value per subject (the surface area of the relevant ROI) that is compared across twin pairs, leading to a single *r*ICC value to characterize the entire population of twins. In contrast, here we measure properties such as cortical curvature that are correlated either spatially across the many vertices of the cortical surface or visually across a grid of points in the visual field, yielding a separate *r* value for each twin pair. We can then compare the distribution of these *r* values for all MZ twins with the distribution for all DZ twins and that for all unrelated pairs. In order to compare data at corresponding cortical locations, the cortical surfaces of subjects were anatomically aligned to the HCP’s *fs_LR* atlas surface, and calculations were limited to a patch of cortex just large enough to contain V1-V3 in all subjects. For correlations over the visual field, properties such as curvature were sampled onto a grid of points in the visual field based on pRF parameters.

Critically, because these are vertex-wise correlations within pairs of subjects, the expected value for unrelated pairs is not 0. For example, the calcarine sulcus is always in approximately the same location between subjects once the surfaces are aligned. Hence the curvature correlations of unrelated pairs are significantly positive. The same patterns hold for other cortical and visual properties. This contrasts with the size correlations across pairs of subjects (previous section), in which the expected value for unrelated pairs is 0.

The properties we chose to compare were the PRF center coordinates *x* and *y*, both expressed in degrees of the visual field, the cortical curvature, and the cortical magnification calculated from the PRF scans. The curvature (**Fig. 10**, left column) and V1 cortical magnification (**Fig. 10**, right column) were calculated both over the cortical surface (top row) and the visual field (bottom row). The PRF *x* and *y* coordinates, which describe the relationship between the cortex and the visual field, appear in the middle column of **Figure 10**. The V1 cortical magnification, was *z*-scored across subjects at each position in the visual field (**Fig. 10F**); without this normalization, the correlations are extremely high for even unrelated pairs. In panel C, correlations were calculated over the shared points in any two subjects’ V1s; however, because V1 does not consist of an identical set of points across subjects, the *z*-score normalization was not possible for panel C.

**Figure 10.**
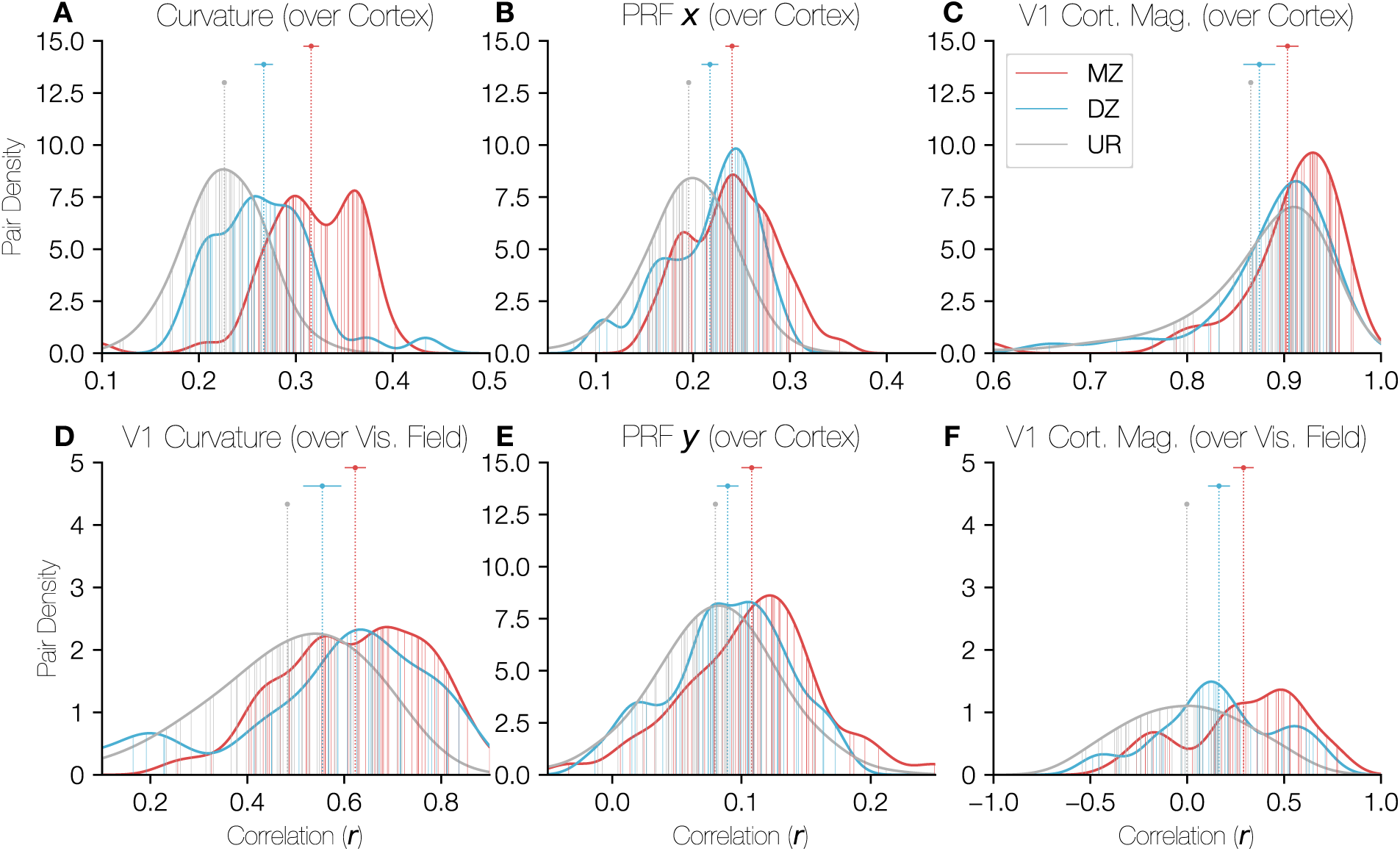
Distributions of correlations of cortical properties between twin pairs. In each panel, the density of correlation values is plotted for all three relationship types (MZ, DZ, UR/unrelated) with the *x*-axis corresponding to the correlation value of the relevant property over the occipital cortical surface and the *y*-axis corresponding to the density of twin pairs at that correlation value. The ROI over which the correlation was calculated was identical for panels A, B, and E; this region was defined on the HCP’s fs_LR anatomically-aligned surface and consisted of a circular region of cortex just large enough to contain V1, V2, and V3 for all subjects. Panel C was restricted to V1 only, while panels D and F were restricted to V1, but correlations were calculated over the visual field rather than over the cortical surface. The points and error-bars at the top of the panels given the median and the 68% confidence interval of the correlation values for each relationship type. Note that in all cases, the unrelated pairs are so numerous that the confidence interval is narrower than the plot-point. Vertical lines plotted from the *x*-axis to the histogram boundary indicate the individual twin-pairs for MZ and DZ groups. For the unrelated pairs (UR), the lines are too numerous to plot, so a random sample of 50 unrelated pairs are shown (however the UR histograms are calculated using all unrelated pairs).

All properties show a pattern whereby the correlations between MZ pairs are significantly higher than those for DZ pairs, which in turn are higher than those for unrelated pairs. Notably, in each case the 68% confidence intervals between MZ/DZ, MZ/UR, and DZ/UR pairs are non-overlapping with the only exception being the DZ/UR overlap in panel C. These results imply that both the topographical distribution of these functional measures on cortex and their distribution over the visual field is influenced by genetic factors. The correlations for **Figures 10A**, **10B**, **10C**, and **10E** are computed after registration to the FreeSurfer template, a process guided by the cortical curvature pattern. Therefore, these correlations likely reflect similarity in both the shape of cortex and the way in which these features are linked to the shape. However, the correlations in **Figures 10D** and **10F** were calculated over the visual field using the functional PRF parameters and thus have no dependency on cortical alignment. These results complement the size correlations reported in the previous section: the size correlations show a similarity in the overall size of visual areas between twins, and these results show similarity in the organization and structure within the maps.

## Discussion

This paper describes an examination of the precise functional boundaries of the early visual cortex in a pool of 181 subjects from the HCP that is the largest of its kind to date. Because our dataset contains V1-V3 visual area boundaries drawn by four trained individuals, it is valuable both for the examination of early visual organization across subjects as well as for understanding the variations across individuals in how such boundaries are drawn. Although similar efforts to characterize the organization of early visual areas have been made by previous researchers (Benson et al., 2014; Wang et al., 2015), our dataset differs in four critical ways. First, our dataset is larger than any similar dataset and consists entirely of data collected at 7T. The large size is especially valuable when trying to estimate variability across the population. Accurate estimates of variability are important for establishing whether newly measured individuals or groups fall within a normal range. Second, the annotations we provide include not just the visual area boundaries but also iso-eccentricity contours that annotate valuable information about the organization of retinotopic maps within each visual area. The various contours can be used to quantify the cortical magnification function as well as the similarities between subregions of the maps. Third, our dataset contains a large number of twin pairs. The combination of variability among unrelated individuals and the similarity between twin pairs sheds light on extent to which the visual organization of cortex is driven by genetic or environmental factors. Finally, our entire dataset, from the pixels originally marked by the anatomists to the figures in this paper, is publicly available and can be easily downloaded and examined by other researchers (https://osf.io/gqnp8/).

### The Value of Functionally Defined Visually Field Maps in Individual Subjects

A great deal of human brain research is conducted by analyzing data in groups of subjects aligned to a template brain, such as the MNI or Talairach volumes (Talairach and Tournoux, 1988; Mazziotta et al., 2001) or the FreeSurfer *fsaverage* surface (Fischl et al., 1999). Alignment to a template is a core step in statistical parametric mapping, perhaps the most widespread approach to analyzing fMRI data (Friston, 2007). This method can be used to predict ROIs (Hinds et al., 2008; Benson et al., 2014; Wang et al., 2015); however it assumes a functional correspondence in anatomically aligned brain locations, an assumption that may often fail (Haxby et al., 2011), leading to high variability in the quality of predictions across subjects (Benson and Winawer, 2018). An alternative approach is to identify specific regions using function responses or a combination of anatomical landmarks and functional responses in individual subjects (Saxe et al., 2006; Benson et al., 2014; Benson and Winawer, 2018). We have taken this latter approach here, annotating individual visual maps by hand. An important observation is that in many cases, alignment by anatomy alone (surface curvature) leads to results that are obviously incorrect relative to the functionally defined areas, in the extreme cases with no overlap at all between anatomically drawn and hand-drawn boundaries. **Figure 3** confirms this variability. In the top panels, the polar angle maps of HCP subject 221319 are shown along with the boundaries of both the anatomically-defined Wang-2015 atlas (white) and the anatomist-drawn contours (black) while in the bottom panel shows a similar plot for subject 111312. The agreement between the hand-drawn boundaries and the anatomically-defined boundaries is low for subject 221319 but high for subject 111312. The hand-drawn contours (black) are also a better match to the polar angle reversals for subject 221319 than the anatomically-defined contours. This demonstrates that the hand-drawn contours are capturing additional functional information that is not captured by the folding pattern alone. That this is true for even early sensory areas suggests that purely anatomical alignment for cortex is likely to risk poor alignment of function for many regions.

Incorporating functional data into the definition of regions of interest does not necessarily entail manual delineation. The manual approach we took here is time-consuming and requires substantial training and validation. It would therefore be useful if the process could be automated. An explicit goal in creating this dataset was to establish a gold-standard set of annotations about the visual cortex in individual subjects against which other models and techniques can be validated and trained. We describe some of these methods below, but note that they were completed without the use of our large, hand-labeled gold standard data set for training.

One method to improve the boundary predictions of anatomical atlases is to combine the atlas prediction (as a prior) with retinotopic measurements (an observation) in a fundamentally Bayesian way (Benson and Winawer, 2018). However, while this method captures individual variation, it is limited by the accuracy of the underlying prior model. Improving this prior model, circularly, requires well-characterized visual areas to study. Additionally, without a dataset annotating the ground-truth of the V1-V3 boundaries, evaluating the accuracy of such predictions in a quantitative way is difficult. Other recent work has begun to examine the application of convolutional neural networks for predicting retinotopic parameters and maps (Thielen et al., 2019; Ribeiro et al., 2020). The dataset provided here is valuable as a potential training and validation resource for future machine learning research on visual cortex. However, a dataset such as this one is only as valuable as it is reliable. One metric for this reliability is the inter-rater agreement across anatomists. To this end, we have provided both the raw data from each anatomist as well as summary data averaged across anatomists for further study. The variability between anatomists in our dataset is small—in particular, the disagreement between anatomists, both across the entire subject population as a whole and for any given subject is minor relative to the differences in retinotopic organizations between subjects. To facilitate usage by other group, all preprocessed and projected data were saved in common open file formats (JSON, HDF5) with wide language support and were uploaded to the OSF (https://osf.io/gqnp8/; DOI: 10.17605/OSF.IO/GQNP8). All source code used to preprocess the contours is open source and can be found on the OSF site. The dataset, including the unprocessed source data from the anatomists and all processed data analyzed in this paper, is also included in the Neuropythy library as a native dataset (https://github.com/noahbenson/neuropythy).

### Enormous Variability in the Size of Early Visual Field Maps

One of the explicit advantages of the dataset described here is its size: to date, no other public dataset of retinotopic maps has a comparable number of subjects as the HCP retinotopy dataset. This project expands the value of the existing retinotopy dataset by also providing visual area boundary annotations. These annotations have allowed us to quantify the distribution of visual area sizes in the population at a previously impossible resolution.

One of the striking features of this distribution is its range: the smallest and the largest instances of each visual area differ by a factor of three or more. This factor is larger than that found in previous work. An examination of 52 cadaverous human hemispheres using the stria of Gennari found this factor to be to be ∼2.9 (Stensaas et al., 1974), while a previous study of 14 hemispheres using functional boundaries between 2-12° of eccentricity found this factor to be ∼2.4 (Dougherty et al., 2003). Similar work in the macaque has found a smaller ratio of ∼2.2 (Van Essen et al., 1984). The ratio of largest to smallest is sample-size-dependent. For example, with a coefficient of variation of 0.2, one expects to find about a 3.3-fold range with 180 measurements (similar to our measures), but only a 2.2-fold range with about 20 measurements (similar to prior measures). Although previous work has found evidence that the size of V1 can weakly predict Vernier acuity (Duncan and Boynton, 2003) as well as the strength of certain optical illusions (Schwarzkopf et al., 2011), these studies do not demonstrate anything like a 3.5-fold (or even 2.2-fold) difference in behavioral effects. Notably, the surface area of the cortex as a whole has a difference of only approximately 1.5-fold between extrema, indicating that the size of V1 is not simply scaled up or down with the size of cortex. In fact, when normalized by the overall cortical surface area, the distribution of relative V1 sizes strongly resembles that of the overall V1 sizes. The correlation between V1 size and cortex size in the dataset is significant but low: *r* = 0.35. However, some of this correlation can be accounted for by the difference in cortex size between males and females; for males only and females only, the correlation is 0.32 and 0.23, respectively. This leaves open the question of why some V1s are so much larger than others and leaves open the possibility that experience plays a substantial role in visual area organization.

Despite low correlations between cortical surface area and V1 surface area, the correlations between V1 and V2, V2 and V3, and V1 and V3 surface areas are high, as are correlations between matched LH and RH visual area sizes and between ventral and dorsal sections of each visual area, indicating that, although the visual area size may not depend strongly on overall cortex size, there is nonetheless some overall shared variability across the early visual system. This observation is consistent with previous results showing that the sizes of the LGN, the optic tract, and V1 vary by a 2- to 3-fold factor across brains but are all strongly correlated within a brain (Andrews et al., 1997), as well as with previous calculations of correlation between V1-V2, V2-V3, and V1-V3 surface area in a small number of subjects (Dougherty et al., 2003). Similar correlations between V1 and V2 have also been reported in the macaque (Sincich et al., 2003). Our dataset demonstrates that this correlation persists beyond V1 in humans to higher order brain areas and provides further evidence that the visual system likely develops in an interdependent manner.

Another intriguing observation from this dataset is the distribution of male and female cortex and V1 sizes. Although the difference is small, the overall size of the cortex of males is larger on average than that of females, and this difference applies to a similar degree to the size of V1, V2, and V3. However, when one normalizes the visual area size by the size of the overall cortex, the distribution of relative visual area sizes for men and women is very similar. This suggests that the differences in visual area size between males and females is a product of the difference in brain size rather than a difference in experience.

A caveat concerning variability is that we compute surface area over only a portion of each map, limited by eccentricity. It is logically possible that the variability would be lower if computed over entire maps (*i.e.*, not limited by 7° of eccentricity). While we don’t have functional definitions of the entire maps, we examined this possibility by computing the variability in size using predictions of the entire visual area sizes as defined using both anatomical alignment (Benson et al., 2014) and extrapolation from the cortical magnification function of Horton and Hoyt (1991). Detailed reports of these calculations both in PDF format (as in **Tab. 2**) and CSV format are available on the OSF repository. While the variability of the anatomically-defined sizes was slightly smaller than that of the measured sizes (CoVs of 0.08-0.11 for V1, V2, and V3), this was true whether the anatomically-defined ROI was limited to 7° or to 90° of eccentricity. Simultaneously, predicted visual area sizes using extrapolation from each ROI’s cortical magnification function had variabilities similar to those of the measured (and 7°-limited) ROIs (CoVs of 0.16-0.22 for extrapolated V1, V2, and V3). The ratios of max:min V1 size for the anatomically-defined and extrapolated ROIs were 2.2 and 3.6, respectively. Although these suggest that our measurements are probably accurate assessments of the entire visual area sizes, it would nonetheless be useful in future work to functionally measure the maps out to a much wider extent in the periphery to verify these estimates.

### Measurements of Cortical Magnification Confirm and Extend Previous Reports

The annotation of both visual area boundaries as well as the boundaries of several sectors of the visual field provides a unique opportunity to measure cortical magnification of V1-V3 in a large population. We find that these measurements, whether calculated using the sector boundaries or using a moving band delimited by iso-eccentricity contours in the visual field, are highly consistent with past reports on average. The mean cortical magnification across subjects, in fact, matched that reported by Horton and Hoyt (1991) almost exactly. Nonetheless, the variation across subjects was large. The variation across subjects occurs in both the total (integrated) cortical magnification, equivalent to visual area size, but also the distribution, meaning that visual areas of different sizes are not simply scaled versions of one another. This is at least qualitatively similar to patterns found in cone density, in which individuals with the highest foveal cone density do not necessarily have the highest parafoveal cone density (Curcio et al., 1990).

Across V1, V2, and V3, the slope of the cortical magnification declines somewhat. Aside from the small decrease in size, and thus cortical magnification, from V1 to V3 (**Fig. 5**), the fraction of each area devoted to the fovea increases relative to the whole area. This effect can be seen both by plotting the eccentricity at which each visual area is split into two equal-area halves (**Fig. 6C**) as well as in the cortical magnification plots of **Figure 6B**, where between 0.5° and 1° of eccentricity, the cortical magnification of the V1 fovea is slightly lower than that of V2 and V3. This effect of the V1-V3 foveal confluence was originally observed by Schira *et al*. (2010), who noted that the V1 cortical magnification dipped below that of V2 and V3 around 0.75° of eccentricity.

A critical point regarding this change in the magnification slope between areas is that the visual areas are not simply rescaled versions of each other: the surface area of V2 is not distributed in a way that is proportional to that of V1—V2’s cortical magnification is steeper than that of V1, and V3’s is steeper than that of V2. If the magnification functions differed only by a scale factor, we would expect the eccentricity value that bisects each visual area to be the same; however in V3 this value is ∼1° lower than in V1.

### Left and Right ROIs are More Symmetric than Ventral and Dorsal ROIs

The pattern of symmetry we see in the brain may reflect the patterns of symmetry we see in natural images. Left-right symmetry is a common feature of many important classes of natural images; for example, images of human faces, plants, and animals all typically have high bilateral symmetry, but relatively lower symmetry when comparing the upper to the lower part of an image (Torralba and Oliva, 2003). This left-right symmetry is also reflected in the high correlation of left and right hemisphere visual areas relative to ventral and dorsal regions of the same visual area. This suggests that the processing requirements for the left and right visual fields may be more similar or more interdependent than those of the upper and lower visual fields with respect to V1-V3’s early role in vision. In fact, V1v and V2v are more similar to each other than to their dorsal counterparts (**Figs. 7E** and **7F**). The same holds true for V2v and V3v compared to their dorsal regions. Although the same observation also holds true for the dorsal regions, the effect is substantially smaller.

One possible explanation of the lower correlation for dorsal areas is measurement noise: the dorsal boundaries may simply be harder to draw. There is substantially more disagreement in the contour positions for the dorsal than the ventral boundaries, especially for V3. Inspection of the retinotopic maps across subjects reveals that the dorsal boundaries are ambiguous in a number of ways and thus may either be harder to draw or may be less consistently organized across subjects. One example of these ambiguities is the occasional presence of a representation of the lower vertical meridian (LVM) for V3 growing out of the LVM representation for V1 (Van Essen and Glasser, 2018; their Fig. S2). This representation occurs in a number of subjects in the dataset and may contribute to variance in the surface areas of dorsal V1-V3. Nonetheless, the overall measure of disagreement is relatively low, even in the dorsal areas, and the correlations between dorsal areas and between ventral-dorsal counterparts are robust, even if lower than for ventral areas.

### MZ Twins’ Retinotopic Maps are More Similar Than Those of DZ Twins or Unrelated Pairs

Visual inspection of twin pairs in this dataset yields the clear observation that twins have more similar retinotopic maps than unrelated pairs. Additionally, MZ twins appear to have the most similar maps among pairs in the dataset overall. In fact, for MZ twins, the correlation between the size of their V1s and that of their twins is ∼0.84 (LH and RH combined; **Fig. 9A**, **Tab. 3**). Remarkably, this is slightly (though likely not significantly) higher even than the correlation between LH V1 and RH V1 (*r* ≈ 0.80).

**Table 3.** Correlations between surface areas across subjects.

| Area 1 | Area 2 | Correlation (*r*) |
| --- | --- | --- |
| Bilateral V1 | Bilateral V2 | 0.77 |
| Bilateral V2 | Bilateral V3 | 0.64 |
| Bilateral V1 | Bilateral V3 | 0.45 |
| Left V1 | Right V1 | 0.80 |
| Left V2 | Right V2 | 0.84 |
| Left V3 | Right V3 | 0.77 |
| Bilateral Ventral V1 | Bilateral Dorsal V1 | 0.68 |
| Bilateral Ventral V2 | Bilateral Dorsal V2 | 0.52 |
| Bilateral Ventral V3 | Bilateral Dorsal V3 | 0.50 |

The close relationship between visual area sizes in twin pairs is partly explainable in terms of the closeness of the anatomical shape between the pairs’ cortices. Comparing the overall cortical surface area of MZ and DZ twins partially supports this idea: for MZ twins, *r* ≈ 0.93 and for DZ twins, *r* ≈ 0.73. However, were the similarity between twins driven entirely by cortical anatomy similarity, we would expect that differences between the functionally defined maps and the anatomically defined maps would be uncorrelated in twin pairs. This was not so, as the residual map areas (functionally defined minus anatomically defined) showed robust correlations. Moreover, the residual correlations were about twice as high in MZ twins as in DZ twins, indicating that functional organization *not* explained by the folding pattern is highly heritable. Thus, although anatomy is highly correlated between twins, especially MZ twins, there is additional similarity between the functional measurements not explained by anatomical differences alone.

Additional comparisons between subject pairs reveal the same trend: regardless of the property being compared, MZ twins are more similar than DZ twins, who are more similar than unrelated subject pairs. This was found for both anatomical properties (cortical curvature) and functional properties of the retinotopic maps (PRF *x*, PRF *y*, and cortical magnification). Our analyses of these properties were calculated using just the occipital cortex, but anatomical data show similar trends when calculated over the entire cortex. Significantly higher correlations for MZ twins than for DZ twins indicates that each of the measures is to some degree heritable, whether that correlation was calculated over the cortical surface, thus depending on cortical alignment, or over the visual field, thus depending only on functional data. While it is possible that correlations over the cortical surface are influenced by cortical alignment, a recent study that performed similar calculations found that the amount of deformation required to align the brains of DZ twins was approximately the same as for MZ twins (Alvarez et al., 2021), indicating that this effect is likely small.

The six properties compared in **Figure 10** as well as the ICC values in **Figure 9** paint a clear story about the similarity of visual cortex and visual cortical organization between twins. Overall, MZ twins are consistently more similar across many measurements than DZ twins, and the difference in surface area between MZ twins is significantly smaller than the same difference between DZ twins: DZ twins’ visual areas are ∼0.6 cm^2^ more different in size than MZ twins, indicating an influence of genetics on the visual cortex. However, these results are also highly consistent with the hypothesis that a shared environment yields similar visual cortices. Notably, both MZ and DZ twins share an environment even in the womb where substantial visual development takes place (Rao et al., 2013). This shared environment during development is important for the Equal Environments Assumption (EEA) that is required to draw conclusions from twin studies. Research into this assumption has found that it is not strictly valid—MZ twins have more similar environments than DZ twins (Guo, 2001)—but that the effects of this incorrectness are usually negligible (Felson, 2014). Nonetheless, it is likely that our sample does not contain enough twins to confidently draw conclusions about heritability on any single metric (Derks et al., 2006).

Despite the consistent pattern of correlations we see, calculations of Falconer’s heritability for size of visual areas using this dataset generally fall short of statistical significance (**Figure 9F**). There are two primary reasons for this. First, our dataset of surface areas is small for heritability studies, which can often have many thousands of twin pairs. It is likely that our calculations of heritability are under-powered, even though the correlations for MZ twins and DZ twins separately are robust. Second, there are a number of sources of noise in our dataset, including imaging noise, noise from subject performance on the retinotopic mapping task, and noise from the anatomists’ boundaries. In calculating correlations, on which heritability calculations are based, compounded sources of noise will have a larger effect on more highly correlated data-points (MZ twins) than those that are less highly correlated (DZ twins and unrelated pairs). (In the extreme case, if the correlation coefficient between two sets of measurements is 100%, additional noise added to the measurements can only reduce the correlation, whereas if the correlation coefficient is 0, noise is equally likely to increase or decrease it.) Thus it is reasonably likely that the population differences in correlations between MZ and DZ twins are larger than the measured differences. Overall, the pattern of results, both in the size correlations and in the correlations of map properties suggests that genetics likely play a large role in the organization of the visual cortex.

These findings regarding the heritability of visual cortex confirm reports from another recently published paper (Alvarez et al., 2021), which evaluated an independent sample of approximately half as many twins as used here. Alvarez *et al*. examined the V1-V3 organization of 36 twin pairs and found both greater visual area overlap in MZ than DZ twins and higher correlations of cortical properties (as in **Figs. 10A**, **10B**) for MZ than DZ twins. Further, the correlations calculated from their dataset approximately agree with the correlations we find here, and both analyses suggest that the heritability of the organization of V3 is higher than that of V2.

## CONCLUSION

We have characterized the functionally-defined structure of early visual areas in human cortex with a previously unmatched precision. Across individuals, variation in the surface area of V1, V2, and V3 is much greater than variation in the size of cortex even when V1-V3 surface areas are normalized by total cortical surface area. Simultaneously, visual area sizes and both anatomical and functional properties of the occipital cortex are highly correlated between twins. In all analyses, correlations were strongest for MZ twins, less strong for DZ twins, and least strong for unrelated pairs. This pattern of correlation across many measurements implies that the organization of early visual cortex is strongly heritable.

